# Unveiling the biochemical catalysis and regulatory mechanisms of the E3-independent E2 enzyme hUBE2O

**DOI:** 10.1101/2025.06.27.661974

**Authors:** Dan Xiang, Xiaoxiao Tang, Ruona Shi, Shuqi Dong, Xiaofei Zhang

## Abstract

UBE2O, an E3-independent E2 ubiquitin-conjugation enzyme, directly engages substrates to mediate Ubiquitin conjugation and ligation. Despite its critical role in ubiquitination of multiple substrates, the catalytic and regulatory mechanisms of *Homo sapiens* UBE2O (hUBE2O) remain incompletely understood. Here, combining domain truncation, systematic mutagenesis and well-designed biochemical approaches, we demonstrate that hUBE2O mediates both mono- and polyubiquitination through a unique catalytic architecture. While none cysteines function for hUBE2O’s E3 activity, the coiled-coil (CC) and C-terminal regulatory (CTR) domains maintain catalytic competence, with the N-terminal regions impose activity constraints. Interestingly, hUBE2O activity is refractory to its self-ubiquitination and phosphorylation state. Instead, specific non-cysteine residues (H939, T995, S1042A, S1046A, S1060A and H1130) emerge as critical regulators of substrate selectivity and catalytic optimization. Surprisingly, zinc ions emerge as potent allosteric inhibitors that bind cysteines of hUBE2O, sterically occluding the access of catalytic site C1040 to Ubiquitin. Our findings reveal that hUBE2O-mediated E3-independent ubiquitination is governed by dynamic interdomain cooperation and allosteric modulation, establishing a mechanistic framework for understanding non-canonical ubiquitination and informing the development of targeted hUBE2O modulators.

## Introduction

Post-translational modifications (PTMs) of proteins dynamically modulate protein activity, stability, localization and interaction networks, thereby governing nearly all cellular processes (1–5). Within the PTMs repertoire, ubiquitination stands out for its versatile mechanism that covalently attaches Ubiquitin to substrate proteins (1, 2, 6). This cascade process is typically orchestrated by three types of enzymes: ubiquitin-activating enzymes (E1), ubiquitin-conjugating enzymes (E2), and ubiquitin ligases (E3) (6–8). In the presence of magnesium ions and ATP, E1 activates Ubiquitin and then form a thioester bond between its catalytic cysteine and the C-terminus of Ubiquitin. Ubiquitin is then transferred to the catalytic cysteine of an E2 enzyme, forming an E2∼Ubiquitin thioester adduct (“∼” represents thioester bond throughout). E3 ligases facilitate substrate specificity by binding both the E2∼Ubiquitin complex and the target protein, catalyzing formation of an isopeptide bond between Ubiquitin and the substrate (6). E3 ubiquitin ligases are classified into three distinct families based on their catalytic mechanisms: RING (really interesting new gene), HECT (homologous to E6AP C-terminus) and RBR (RING-between-RING). RING E3s directly transfer Ubiquitin from E2∼Ubiquitin to substrates. HECT E3s first form an E3∼Ubiquitin intermediate via their catalytic cysteine before transferring Ubiquitin to substrates, whereas RBR E3 utilize a bipartite mechanism combining RING-mediated E2∼Ubiquitin recruitment and HECT-like thioester intermediate formation (8, 9).

While the majority of E2 enzymes strictly depend on E3 partners for functionality, the ubiquitin-conjugating enzyme E2O (UBE2O) stands as one of the only two known E2 enzymes capable of directly binding and catalyzing substrate ubiquitination independently of E3 ligases (10, 11). Biologically, hUBE2O catalyzes ubiquitination of diverse substrates, including SMAD6, BAP1, RECQL4, CTNNA1 (our manuscript under revision in EMBO Reports) and ribosomal proteins, establishing it in critical pathways such as orphan protein quality control, proteome remodeling, DNA repair, cell adhesion and cancer pathogenesis (3, 5, 12–17). Mechanistically, however, the molecular basis for hUBE2O’s autonomous activity and regulatory control remains incompletely understood.

hUBE2O comprises five domains: the N-terminal conserved domains 1–2 (CR1-CR2), coiled coil (CC), the ubiquitin conjugating (UBC) domain, and the C-terminal regulatory (CTR) domain, which contains a C-terminal conserved region 3 (CR3). While CR1 and CR2 domains mediate substrate and adaptor binding (3, 14, 18), the UBC domain contains a catalytic cysteine residue C1040 for Ubiquitin conjugation. However, whether additional catalytic cysteines contribute to its E3 activity, and the functions of the non-catalytic CC and CTR domains, remain unclear. Despite its involvement in multiple biological processes, the regulatory mechanisms governing hUBE2O activity are poorly understood, limiting therapeutic exploitation. Cryo-EM structures of hUBE2O complexed with a cofactor NAP1L1 revealed that the substrate selection of hUBE2O is modulated by Ubiquitin binding and NAP1L1 (18). However, how substrate specificity is achieved in the absence of cofactors remain unsolved. Additionally, while phosphorylation activates the RING E3 CBL (19), whether PTMs or non-protein molecules regulate hUBE2O’s activity requires investigation. Resolving these questions is essential to deciphering hUBE2O’s enzymatic logic.

Here, combining domain truncation, systematic mutagenesis, and AlphaFold3 modeling, we delineate that hUBE2O achieves E3-independent ubiquitination via dynamic interdomain cooperation. Notably, E3 activity of hUBE2O does not require additional catalytic cysteine. Instead, the CC/CTR domains maintain catalytic competence, while CR1-CR2 regions exert activity restrictions. In addition, several polar amino acids in the vicinity of the E2 active site are critical for catalytic activity and substrate specificity of hUBE2O. Counterintuitively, hUBE2O activity is insensitive to phosphorylation or self-ubiquitination but highly responsive to zinc ions. Unlike zinc-activated E3-RING ligases (9, 20), hUBE2O undergoes allosteric inhibition via zinc-mediated occlusion of the catalytic pocket through cysteine residues. These findings provide mechanistic insights into hUBE2O’s unique enzymatic logic and facilitate the development of targeted modulators.

## Results

### hUBE2O functions as an E3-independent E2 enzyme mediating mono- and polyubiquitination

To characterize the enzymatic activity of hUBE2O, we expressed recombinant hUBE2O in Expi293F cells and purified it for in vitro ubiquitination assays. As depicted in Fig. 1A, hUBE2O exhibited robust E3-independent self-ubiquitination activity, manifested as either multi-monoubiquitination or polyubiquitination. Using a lysine-deficient (K0) Ubiquitin mutant to preclude polyubiquitin chain formation, we unambiguously demonstrated that hUBE2O mediates multi-monoubiquitination of itself (Fig. 1A). To investigate potential polyubiquitin chain formation activity, we excised gel slices of the in vitro hUBE2O ubiquitination reaction mixture (Fig. S1A), performed in-gel digestion, and subject the resulting peptides to mass spectrometry analysis. Beyond its established capcity to form K48 linkages (21, 22), hUBE2O generated K63- and K11-linked polyubiquitin chains in vitro (Fig. 1B). Conversely, substitution of wild-type Ubiquitin with K0 Ubiquitin abolished formation of these linkages (Fig. 1C).

**Figure 1.**
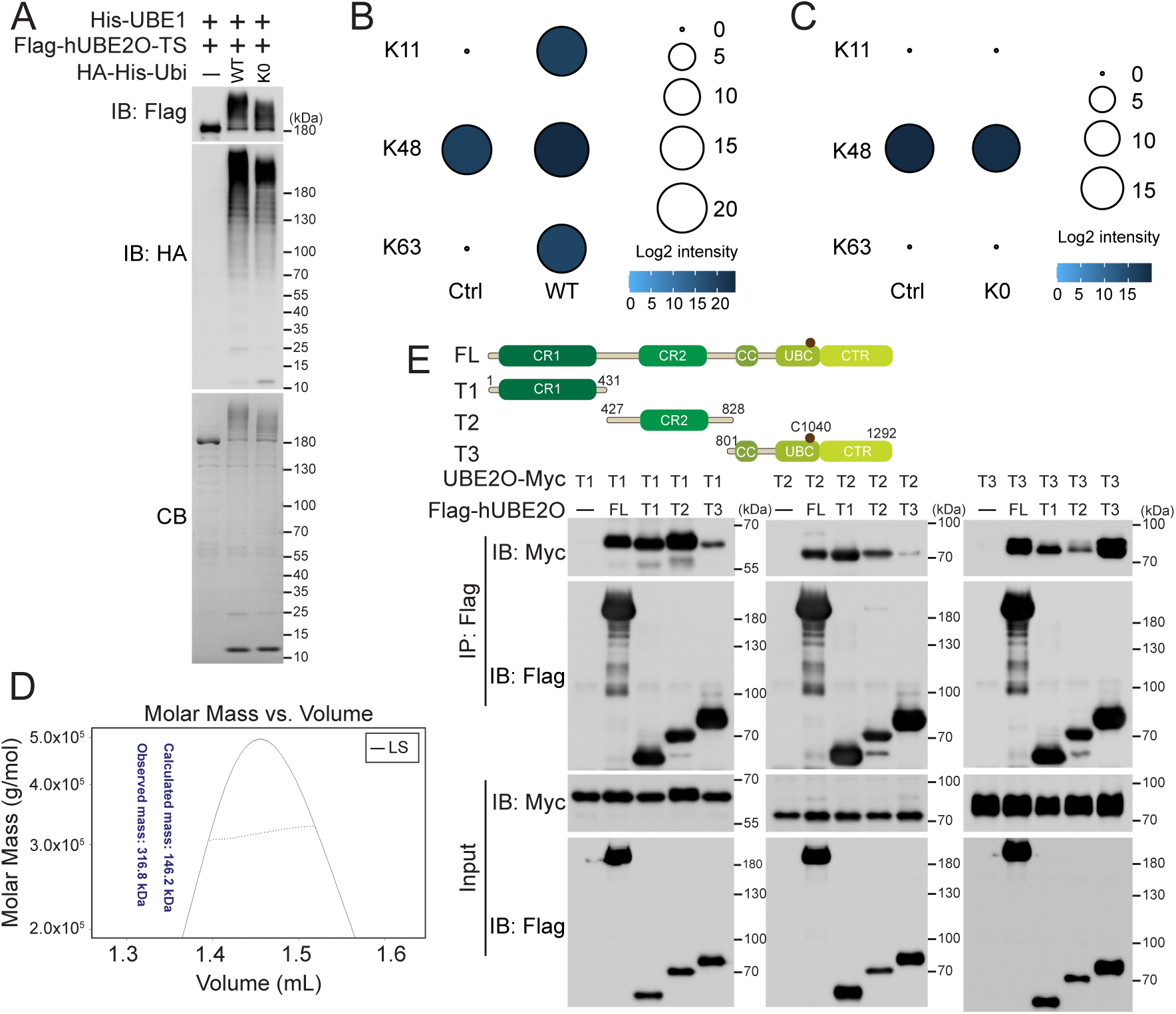
hUBE2O functions as an E3-independent E2 enzyme mediating mono- and polyubiquitination. *A, in vitro* ubiquitination assay shows hUBE2O catalyzes ubiquitination in an E3-independent manner. K0: lysine deficient mutant; IB: immunoblotting; CB: Coomassie blue staining. *B and C,* bubble plots show the log2 intensity of hUBE2O catalyzed polyubiquitin chain peptides identified by mass spectrometry. Detailed data are available in datasets S1. *D,* molar mass analysis of purified Flag-UBE2O-TwinStrep proteins by SEC-MALS. *E,* immunoprecipitation assay shows the interactions between hUBE2O full-length (FL) and its truncations. The top panel shows the schematic diagram of the full-length and truncations of hUBE2O, the E2 catalytic active site (C1040) is indicated by a brown dot. Ubiquitination and interaction experiments were repeated at least twice, one representative result is shown. Source data for this figure are available in supporting information as: SourceDataF1.

During purification processes, size exclusion chromatography (SEC) indicated that hUBE2O eluted as a dimer, a finding was confirmed by SEC-MALS (Size Exclusion Chromatography–Multi-Angle Light Scattering) (Fig. 1D). To delineate interaction domains, we generated truncated hUBE2O variants (Fig. 1E, top panel) and performed co-immunoprecipitation experiments. All three hUBE2O truncations demonstrated interaction with full-length hUBE2O (FL-hUBE2O). Specifically, the CR1 domain preferentially interacted with CR2 domain, whereas the T3 truncation (containing CC, UBC, and CTR domains) exhibited enhanced self-association (Fig. 1E, bottom panel). In summary, these data establish hUBE2O dimerizes through multi-domains interactions and functions as an E3-independent E2 enzyme mediating both mono- and polyubiquitination.

### C1040 is the sole catalytic cysteine residue in hUBE2O

The pioneering work by Berleth and Pickart proposed a mode wherein hUBE2O employs a cysteine residue as an E3-like active site (Cys^E3^) in addition to its canonical E2 catalytic site (11). To clarify whether hUBE2O indeed harbors a cysteine residue that functions as an active E3 site, we systematically mutated each of the 25 cysteines in hUBE2O to serine residues (Fig. 2A, top panel). We expressed these Myc-tagged mutants in HEK293T cells, enriched them using anti-Myc beads, and subsequently performed an in vitro ubiquitination assay. To investigate the impact of cysteine mutations on hUBE2O’s substrate ubiquitination activity, we incorporated a recently identified hUBE2O substrate CTNNA1 into the reaction. We used an anti-V5 antibody to access CTNNA1 ubiquitination, an anti-Myc antibody to evaluate hUBE2O self-ubiquitination and an anti-HA antibody to monitor polyubiquitin chain formation. Strikingly, only the C1040S mutation fully abrogated hUBE2O ubiquitination activities, encompassing substrate (CTNNA1) ubiquitination, self-ubiquitination, and polyubiquitin chain formation (Fig. 2A). While the adjacent C1025S displayed very mild reduced activity relative to wild-type (WT) hUBE2O, evolutionary conservation analysis revealed poor sequence conservation of C1025 across species (Fig. S1B), implicating its potential role in structural support rather than direct catalysis.

**Figure 2.**
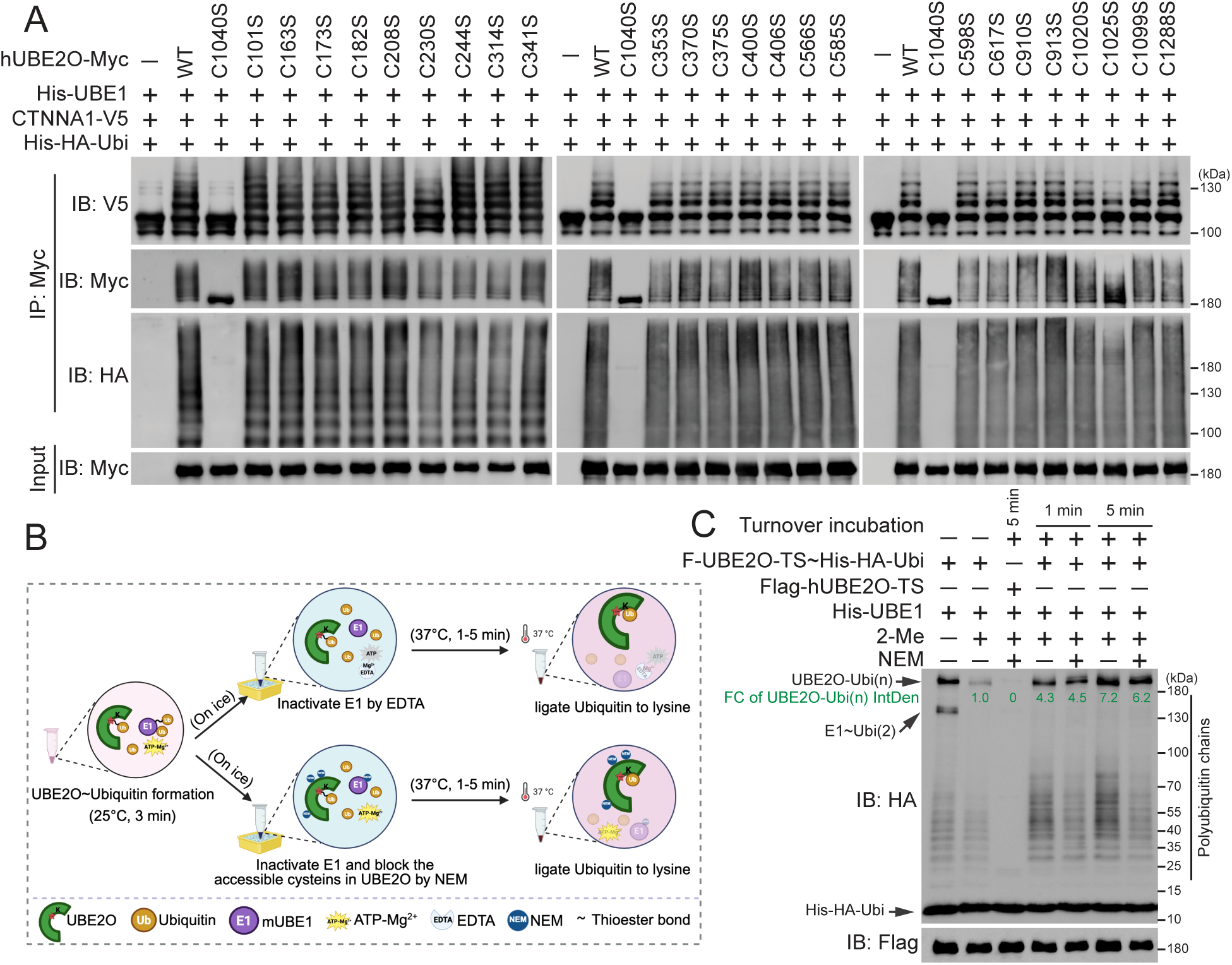
C1040 is the sole catalytic cysteine of hUBE2O. *A,* immunoblots show the self-ubiquitination (IB: Myc), polyubiquitin chain formation (IB: HA) and ubiquitination of CTNNA1 (IB: V5) catalyzed by each of the 25 hUBE2O cysteine to serine mutations in vitro. *B and C,* C1040 is the sole catalytic cysteine of hUBE2O. *B,* schematic diagram showing the assay of hUBE2O∼Ubiquitin adducts formation and turnover. Detailed description is shown in the experimental procedures. This diagram was created in BioRender (https://BioRender.com). *C,* immunoblots show the NEM-blocked hUBE2O∼Ubiquitin adducts turnover ability, the detailed experimental procedures are described in the experimental procedures. All experiments were repeated at least twice, one representative result is shown. Source data for this figure are available in supporting information as: SourceDataF2.

To further dissect the mechanistic roles of cysteine functions, we developed a thioester-trapping strategy. As illustrated in Fig. 2B, brief (3 min) in vitro reaction at room temperature enabled the formation of a thioester bond linked hUBE2O∼Ubiquitin adducts between C1040 and ubiquitin. These adducts are reversible upon treatment with the reductant 2-Mercaptoethanol (Fig. 2C, lanes 1 and 2). The reaction was then rapidly quenched by reducing the temperature (< 4°C), inactivating E1 by EDTA (chelating magnesium ions) or N-Ethylmaleimide (NEM, a thiol-alkylating agent). Notably, NEM can also irreversibly block other accessible cysteines in hUBE2O. Subsequent time-course incubation at 37°C allowed the Ubiquitin moiety of the hUBE2O∼Ubiquitin adducts to transfer to lysine residues (Fig. 2C, lanes 2, 4 and 6). As anticipated, pretreatment with NEM completely abolished hUBE2O enzymatic activity (Fig. 2C, lane 3). Strikingly, NEM treatment after thioester formation did not impair Ubiquitin transfer from E2-active site to lysine residues (Fig. 2C, lanes 2, 5 and 7). These results conclusively demonstrate that cysteines are dispensable for Ubiquitin transfer subsequent to E2∼Ubiquitin formation, indicating that hUBE2O does not require a Cys^E3^.

In the hUBE2O catalytic mechanism proposed by Berleth and Pickart, phenylarsenoxides (PAO) inhibited hUBE2O by binding Cys^E3^ and a vicinal cysteine (11). Cysteine mutagenesis screening revealed that among all 25 cysteine-to-serine mutations of hUBE2O, the vicinal C910S and C913S mutations conferred resistance to PAO mediated inhibition (Fig. S1C). However, both C910S and C913S mutations maintained full E3 enzymatic activity (Fig. 2A), indicating the PAO-mediated hUBE2O inhibition is not through E3 site occupation. Although a prior work showed hUBE2O mediates K48 polyubiquitination of BMAL1 and identified C617 of hUBE2O as the active E3 site in this process (23), we were unable to replicate ubiquitination of BMAL1 by hUBE2O or observe the E3 activity of hUBE2O on BMAL1 (Fig. S1, D and E). Furthermore, our data showed that C617S mutant retained fully activity for hUBE2O-mediated CTNNA1 ubiquitination and self-ubiquitination activity (Fig. 2A). Collectively, our comprehensive mutagenesis and biochemical analyses demonstrate that C1040 is the only catalytically essential cysteine residue in hUBE2O.

### Roles of hUBE2O domains in its ubiquitination activity

To further delineate the structural determinants of hUBE2O’s enzymatic activity, we generated a series of domain deletion mutants (Fig. 3A). Following expression in HEK293T cells and anti-Flag affinity purification, each mutant was tested for self-ubiquitination, polyubiquitin chain formation and CTNNA1 ubiquitination activities. Mutants lacking CR1, CR2 or CC domains (D1-D3) retained full self-ubiquitination and polyubiquitin chain formation activities (Fig. 3B). In contrast, mutants lacking the CTR domain (D4-D7) completely lost these activities, as well as the ability to ubiquitinate CTNNA1 (Fig. 3, B and C). Truncations retaining the CTR domain but lacking the CC domain (D3, D8) strongly impaired CTNNA1 ubiquitination (Fig. 3C and D), consistent with previous reports using SMAD6 as a substrate of hUBE2O (14). Importantly, the UBC domain alone (D5) or truncations without CTR domain (D4, D6) were unable to form the UBC∼Ubiquitin thioester intermediate (Fig. 3E), indicating the CTR domain is essential for hUBE2O to execute its E2-conjugationg activity. To identify the minimal functional elements, we generated additional truncations within the CTR domain. The D10 mutant, which lacks flexible regions unresolved in the hUBE2O-NAP1L1 complex structure (PDB:7UN6), exhibited reduced activities compared to D9 (Fig. 3F). Further truncation by one amino acid deletion in D10 (D11) resulted in a significant comprise of the CTNNA1 ubiquitination (Fig. 3F). Intriguingly, both D10 and D11 retained self-ubiquitination ability, albeit with less activity compared to D9, indicating a dependence on the structural flexibility. These results demonstrate that while the UBC domain contains the catalytic core C1040, the CTR domain is indispensable for the transfer of Ubiquitin from E1∼Ubiquitin to the C1040 of UBC. In addition, CC domain is requisite for the maximal substrate-directed activity of hUBE2O.

**Figure 3.**
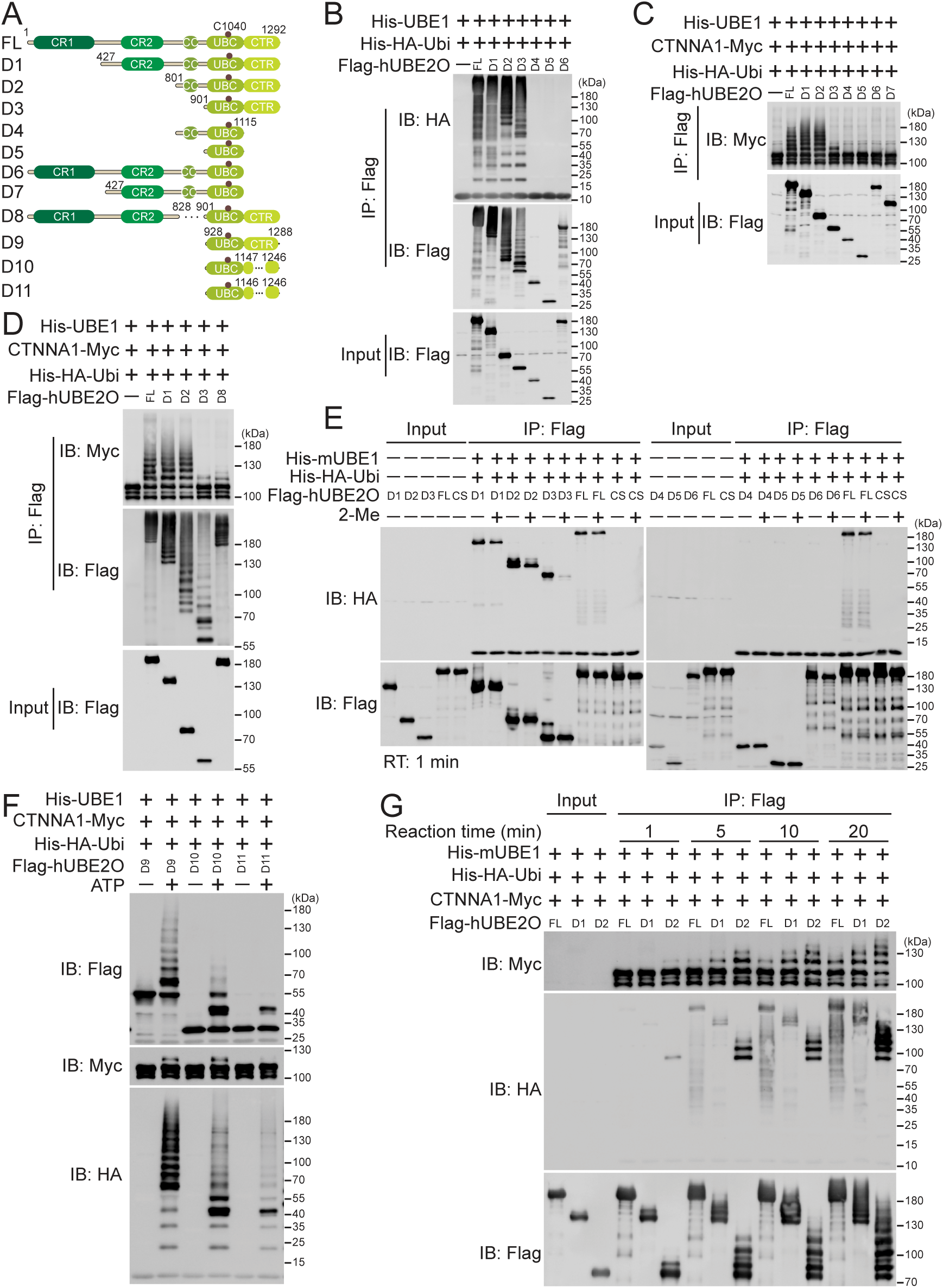
Roles of domains of hUBE2O in ubiquitination. *A,* schematic diagram of the full-length and deletions of hUBE2O, the E2 catalytic active site (C1040) is indicated by a brown dot. *B,* in vitro ubiquitination assay shows the self-ubiquitination (IB: Flag) and polyubiquitin chain formation (IB: HA) abilities of the indicated hUBE2O deletions expressed in HEK293T cells. *C* and *D,* in vitro ubiquitination assay shows the CTNNA1 ubiquitination catalyzing ability (IB: Myc) of the indicated hUBE2O deletions expressed in HEK293T cells. *E,* E2∼Ubiquitin adducts formation assay of the indicated hUBE2O deletions expressed in HEK293T cells. *F,* in vitro ubiquitination assay shows the self-ubiquitination (IB: Flag), polyubiquitin chain formation (IB: HA) and CTNNA1 ubiquitination catalyzing (IB: Myc) abilities of the indicated hUBE2O deletions expressed in HEK293T cells. *G,* immunoblots show the time-course enzymatic activity of the indicated hUBE2O deletions expressed in HEK293T cells. All experiments were repeated at least twice, one representative result is shown. Source data for this figure are available in supporting information as: SourceDataF3.

Interestingly, the D2 mutant exhibited higher self-ubiquitination and substrate ubiquitination activities than FL-hUBE2O (Fig. 3B-D). Time-course experiments demonstrated the fastest accumulation of ubiquitinated products among FL-UBE2O and D1, indicating the enhanced ubiquitination efficiency of D2 (Fig. 3G). These findings suggest that the N-terminal regions (CR1/CR2) exert an inhibitory effect on the ubiquitination activity of the C-terminal regions of hUBE2O.

### Self-ubiquitination exhibits minimal regulation of hUBE2O activity in vitro

To explore potential regulatory mechanisms of hUBE2O activity, we considered multiple factors known to modulate E2-E3 systems, including post-translational modifications, protein-protein interactions, and small molecule binding (24). Our mass spectrometry analysis of self-ubiquitinated hUBE2O identified 27 GlyGly-modified lysine residues in hUBE2O (Figs. S1A, and S2A). Considering the extensive self-ubiquitination of hUBE2O and a recent study showing that self-ubiquitination in the CR1-CR2 region enhances the E3 activity of *Trametes pubescens* (tp) UBE2O (25), we first generated single-point mutations at three key sites: reported cancer-associated mutations K825/K953 (12) and K846, the ubiquitinated site in our MS data between K825 and K953. Unfortunately, none of these individual mutations affected hUBE2O’s activity (Fig. S2B). We therefore generated a lysine-null mutant (K0-UBE2O) by replacing all 87 lysine residues with arginine residues. As expected, K0-UBE2O completely lost self-ubiquitination capacity (Fig. S2C). Strikingly, K0-UBE2O retained full activities for CTNNA1 ubiquitination and polyubiquitin chain formation (Fig. S2C), although the hUBE2O∼Ubiquitin thioester intermediate signal was weaker in K0-UBE2O during short reaction time (3 min) due to the absence of lysine acceptors (Fig. S2D). Taken together, these results demonstrate that, in contrast to tpUBE2O (25), hUBE2O’s catalytic activity is independent of its self-ubiquitination status.

### The enzymatic activity of hUBE2O is restricted by specific residues within the UBC domain

Apart from ubiquitination, previous study has reported that phosphorylation of hUBE2O enhances its interaction with mixed-lineage leukemia (MLL) protein, resulting to MLL polyubiquitination and degradation (26). Leveraging PhosphoSitePlus site (https://www.phosphosite.org/homeAction.action), we identified multiple phosphorylation sites in hUBE2O, suggesting potential regulatory functions of phosphorylation on hUBE2O’s activity. To investigate this possibility, we carried out comprehensive mutations of all serine, threonine, and tyrosine residues within the CC, UBC, and CTR domains (essential for the maximal activity of hUBE2O). Initial screening revealed that the combined mutation of several residues in the UBC domain completely abolished both self-ubiquitination and polyubiquitin chain formation activities (Fig. S3A). Subsequent individual alanine substitution demonstrated that T995A, S1042A, S1046A, and S1060A mutated hUBE2O each resulted in a significant reduction in enzymatic activities when compared to WT-UBE2O in in vitro ubiquitination system (Fig. 4A).

**Figure 4.**
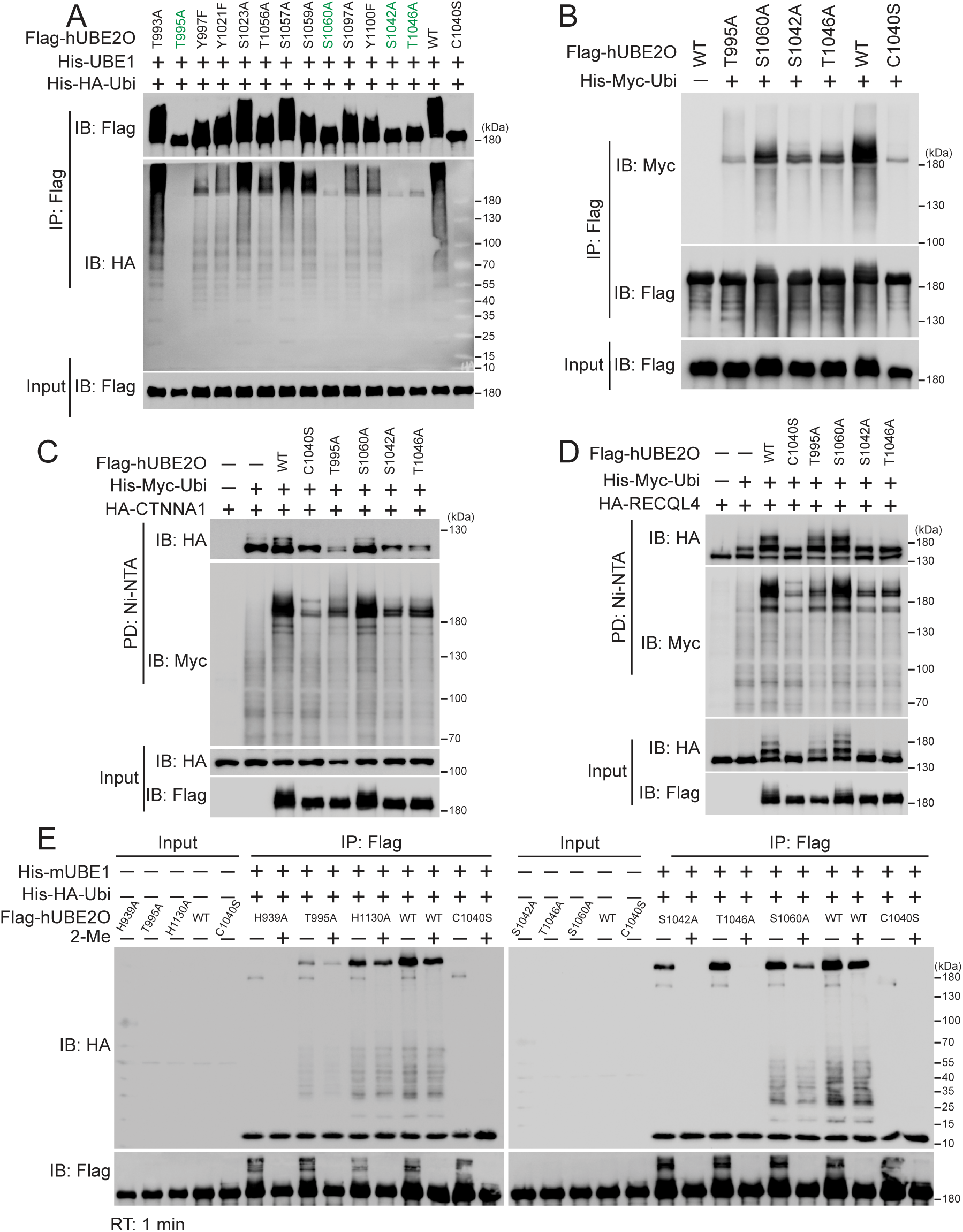
The enzymatic activity of hUBE2O is restricted by specific residues within the UBC domain. *A,* in vitro ubiquitination assay shows the self-ubiquitination (IB: Flag) and polyubiquitin chain formation (IB: HA) abilities of the indicated hUBE2O mutations expressed in HEK293T cells. *B,* immunoblots show the self-ubiquitination activity (IB: Myc) of the indicated hUBE2O mutations in HEK293T cells. *C* and *D,* immunoblots show CTNNA1 (*C*) and RECQL4 (*D*) ubiquitination catalyzing (IB: HA) activities of the indicated hUBE2O mutations in HEK293T cells. *E,* E2∼Ubiquitin adducts formation assay of the indicated hUBE2O mutations expressed in HEK293T cells. All experiments were repeated at least twice, one representative result is shown. Source data for this figure are available in supporting information as: SourceDataF4.

In cellular ubiquitination assays conducted in HEK293T cells, the T995A mutation showed the most pronounced effect, reducing self-ubiquitination to levels comparable to those of the C1040S mutant (Fig. 4B). When assessing substrate specificity using CTNNA1 and RECQL4 (3) as model substrates, we observed that S1042A and S1046A similarly attenuated ubiquitination of both substrates. In contrast, T995A displayed a more selective effect, severely impairing CTNNA1 ubiquitination while only mildly affecting RECQL4 ubiquitination (Fig. 4, C and D). These results suggest that T995 plays a particularly important role in substrate selection.

To further dissect the mechanistic underpinnings of these effects, we performed an in vitro hUBE2O∼Ubiquitin adducts formation assay using these mutants. This analysis revealed that T1046A and S1060A exhibited hUBE2O∼Ubiquitin adducts formation ability indistinguishable from WT-UBE2O, whereas the T995A showed profound impairment in E2 catalytic activity (Fig. 4E). Interestingly, efforts to rescue the activity of these mutants through phosphorylation-mimetic mutations were unsuccessful (Fig. S3B). Conversely, substitution of T995A to serine (T995S) -another polar amino acid-restored enzymatic activity of hUBE2O-T995A (Fig. S3B), indicating that the functional constraints at this position are governed by structure rather than phosphorylation-dependent requirements. Together, these findings demonstrate that hUBE2O’s activity is fine-tuned by specific residues within the UBC domain, with T995 playing a particularly important role in both catalytic activity and substrate specificity.

### Zinc ions inhibit the enzymatic activity of hUBE2O both in vitro and in vivo

To further examine whether phosphorylation status of hUBE2O affects its enzymatic activity, we adopted an alternative experimental strategy. We expressed Flag-tagged hUBE2O in HEK293T cells, immunoprecipitated the protein using anti-Flag beads, and treated the purified protein with either Lambda Protein Phosphatase (Lambda PP) or FastAP Thermosensitive Alkaline Phosphatase (FastAP) to remove phosphate groups. Following extensive washing to remove the phosphatases, we performed in vitro ubiquitination assays with E1 and Ubiquitin. Lambda PP treatment effectively reduced threonine phosphorylation of hUBE2O without compromising its self-ubiquitination or polyubiquitin chain formation capabilities. In contrast, FastAP treatment completely abolished the catalytic activity of hUBE2O (Fig. S4A).

To ensure whether the observed activity block of hUBE2O by FastAP resulted from dephosphorylation or other ingredients in the dephosphorylation reaction mixture, we pre-incubated hUBE2O with heat-inactivated phosphatase, reaction buffer for dephosphorylation reaction or storage buffer for phosphatase prior to ubiquitination reaction. Control experiments revealed that neither active phosphatase Lambda PP nor any of the phosphatase reaction buffers inhibited hUBE2O’s activity. In contrast, pre-incubation of hUBE2O with storage buffer of either FastAP or Quick CIP phosphatase caused inhibition comparable to that of the active enzymes (Fig. S4B). This suggested that the observed inhibition originated not from phosphatase activity but from components in the storage buffer.

Further analysis of the storage buffer ingredients of FastAP and Quick CIP identified zinc ions as the most probable inhibitory component of hUBE2O activity. Dose-response experiments confirmed that zinc chloride inhibited hUBE2O activity in a concentration-dependent manner (Fig. 5A). Using a colorimetric assay, we detected a significant reduction in free zinc ions concentration upon hUBE2O addition, confirming direct binding of hUBE2O and zinc ions (Fig. 5B). Moreover, the inhibition was specific to zinc ions, as neither calcium nor manganese ions affected hUBE2O activity under identical conditions (Fig. 5C).

**Figure 5.**
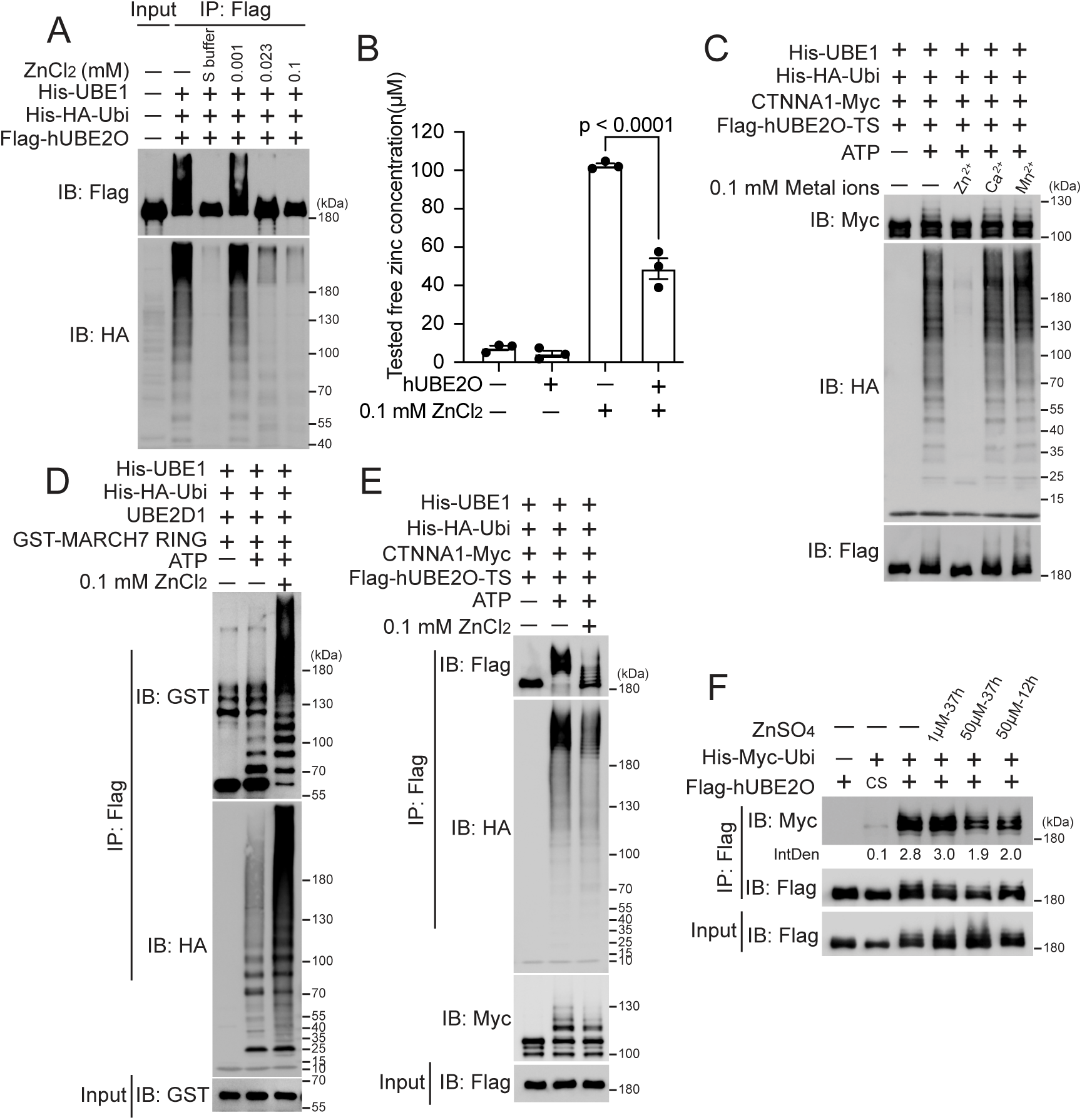
Zinc inhibits the enzymatic activity of hUBE2O both in vitro and in vivo. *A,* immunoblots show zinc ions inhibit the self-ubiquitination (IB: Flag) and polyubiquitin chain formation (IB: HA) activities of HEK293T-expressed hUBE2O in a dose-dependent manner in vitro. *B,* zinc colorimetric assay shows the concentration of free zinc ions in solution significantly decreases upon hUBE2O addition. n = 3 replicates in all cases. Error bars indicate mean ± SEM. Significance among multiple groups was determined using ANOVA followed by Tukey’s post hoc test. *C,* immunoblots show the self-ubiquitination (IB: Flag), polyubiquitin chain formation (IB: HA) and CTNNA1 ubiquitination catalyzing (IB: Myc) abilities of purified hUBE2O are inhibited by zinc ions in vitro, but not by calcium nor manganese ions. *D* and *E,* immunoblots show zinc ions enhance MARCH7 RING-mediated ubiquitination (*D*), while specifically inhibiting hUBE2O mediated ubiquitination (*E*). *F,* ubiquitination assay in HEK293T cells shows zinc ions reduce hUBE2O’s self-ubiquitination in cells. All experiments were repeated at least twice, one representative result is shown. Source data for this figure are available in supporting information as: SourceDataF5.

To address whether residual zinc ions bound to hUBE2O could interfere with other components in the ubiquitination system (such as E1 or Ubiquitin itself), we conducted parallel assays to exam the activity of the MARCH7 RING domain under identical conditions. In line with known fact (9, 20), we observed that zinc ions enhanced MARCH7 RING-mediated ubiquitination while inhibiting hUBE2O-mediated ubiquitination at the same concentration of zinc ions, Ubiquitin and E1 (Fig. 5, D and E). This contrasting effect provides compelling evidence that zinc ions selectively inhibit hUBE2O rather than broadly disrupting the ubiquitination machinery. In addition, zinc ions reduced hUBE2O self-ubiquitination in vivo without affecting its solubility/accumulation or overall polyubiquitination patterns (Figs. 5F and S4C). Notably, copper treatment produced the previously reported increase in polyubiquitination (27). Collectively, these results establish zinc ion as a specific inhibitor of hUBE2O enzymatic activity both in vitro and in cellular contexts.

### Zinc ions inhibit the activity of hUBE2O by hindering the accessibility of Ubiquitin to hUBE2O’s catalytic C1040 residue

To elucidate the structural basis for zinc ion-mediated inhibition of hUBE2O activity, we systematically mapped potential binding sites. We first analyzed structural domains of hUBE2O. Both D2 and D3 deletions remained sensitive to zinc ions inhibition (Fig. 6A), localizing the binding site of zinc ions to the UBC-containing D3 deletion (UBC and CTR domains). Given that zinc ions are known to coordinate with cysteine and histidine residues in E3-RING domains (9, 20), we performed subsequent cysteine mutagenesis screening. As shown in Fig. 6B, C910S, C913S and C1020S mutations conferred resistance to zinc ions inhibition, identifying these three cysteines in UBC domain as critical zinc-binding residues. Interestingly, although only C910 is evolutionarily conserved between *Homo sapiens* and *Danio rerio* (dr, zebrafish) UBE2O (Fig. S1A), zinc ions still inhibited catalytic activity of drUBE2O (Fig. S5A), suggesting broad conservation of this regulatory mechanism across species.

**Figure 6.**
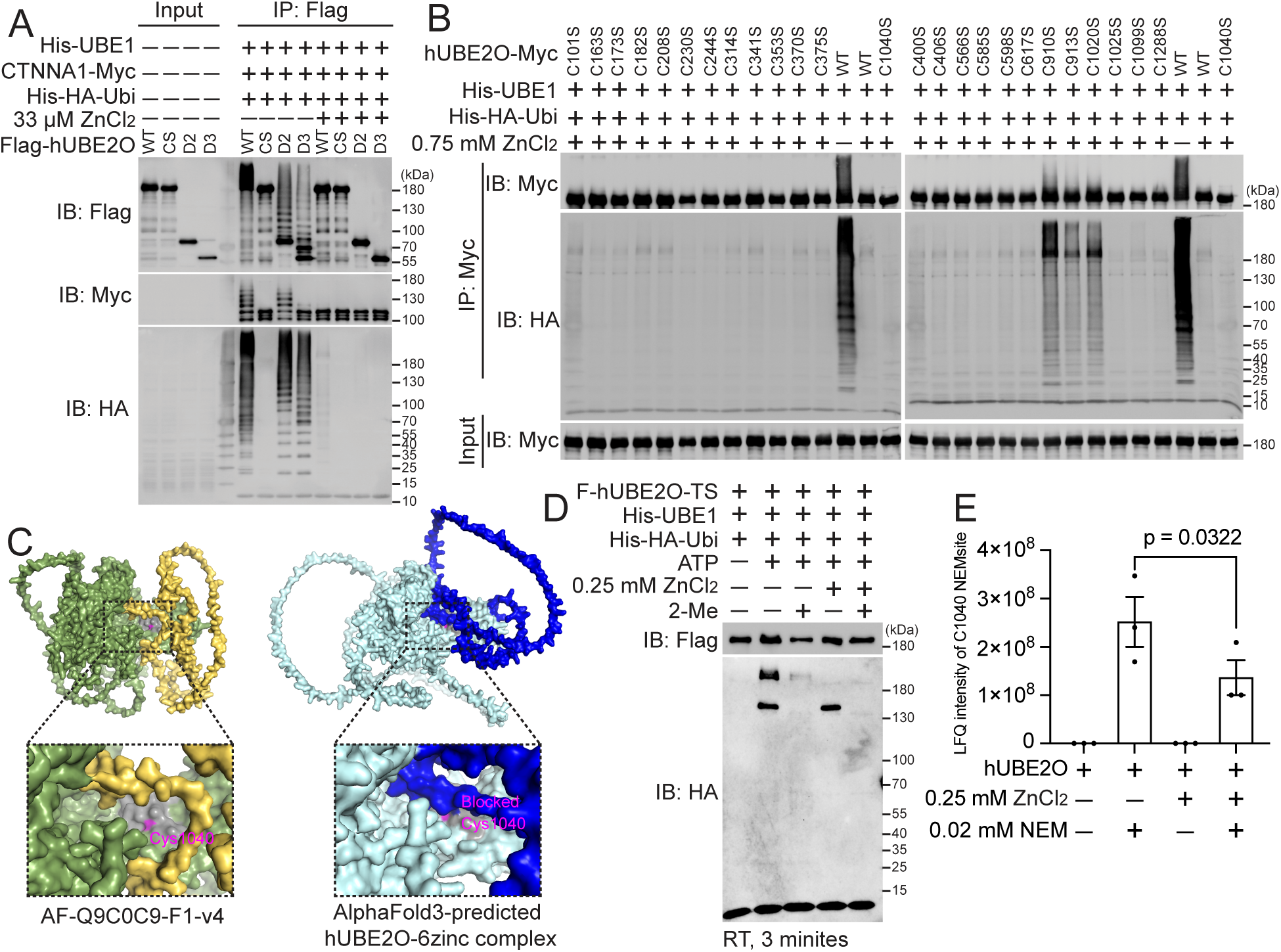
Zinc inhibits the activity of hUBE2O by hindering the accessibility to its catalytic C1040 residue. *A,* immunoblots show the self-ubiquitination (IB: Flag), polyubiquitin chain formation (IB: HA) and CTNNA1 ubiquitination catalyzing (IB: Myc) abilities of HEK293T-expressed hUBE2O deletions with or without the addition of zinc ions. CS: C1040S. *B,* immunoblots show C910S, C913S, and C1020S mutations are resistant to zinc ions-mediated inhibition among all 25 hUBE2O cysteine to serine mutations. *C,* structural models of apo hUBE2O (left) or hUBE2O in complex with six zinc ions (right) generated using AlphaFold3, with zinc ions bound at the C910 and C913 residues (right). Both structures are shown as cartoon representation with individual regions color coded. The C1020 to T1046 residues (gray) and C1040 residue (magenta) of both structures are shown as cartoons. The N709 to W902 residues are shown as yellow cartoon in apo hUBE2O or blue cartoon in hUBE2O-6zinc complex. The rest of regions are shown as green cartoon in apo hUBE2O or palecyan cartoon in hUBE2O-6zinc complex. The catalytic cores of both structures are enlarged and C1040 residue is labeled in magenta (bottom). *D,* E2∼Ubiquitin adduct formation assay shows zinc ions abolish hUBE2O’s ability to form the hUBE2O∼Ubiquitin adducts. *E,* bar plot shows the mass spectrometry-identified label free quantification (LFQ) intensities of NEM-modified C1040 peptides with or without the presence of zinc ions. n = 3 replicates in all cases. Error bars indicate mean ± SEM. Significance among multiple groups was determined using ANOVA followed by Tukey’s post hoc test. Detailed data are available in datasets S1. All experiments except for mass spectrometry identification were repeated at least twice, one representative result is shown. Source data for this figure are available in supporting information as: SourceDataF6.

To further characterize potential zinc ion binding sites, we expanded our mutagenesis analysis to all five histidine residues within the hUBE2O-D3 deletions. Among these, the H939A and H1130A mutations led to the loss of enzymatic activity of hUBE2O (Fig. S5B), thereby precluding an assessment of zinc ion sensitivity. The remaining histidine mutants (H1018A, H1149A and H1215A) retained catalytic function but exhibited no resistance to zinc-mediated inhibition of hUBE2O activity (Fig. S5B), thus excluding these histidine residues as major zinc ions binding sites (Fig. S5B).

To elucidate the structural mechanism of zinc ions-mediated inhibition of hUBE2O activity, we employed AlphaFold3 for computational modeling (28). The predicted structure revealed direct coordination of zinc ions by three cysteine residues (C910, C913 and C1020) within the UBC domain of hUBE2O (Fig. S5C). Comparative analysis with the wild-type apo hUBE2O structure showed that zinc ions binding induced significant steric occlusion of the hUBE2O catalytic pocket (Fig. 6C), providing a structural rationale for the observed inhibition. Biochemical assays confirmed this structural prediction: first, treatment with zinc ions abolished the formation of hUBE2O∼Ubiquitin thioester intermediate (Fig. 6D); second, mass spectrometry analysis revealed that the zinc ions reduced the modification efficiency of the catalytic cysteine (C1040) by NEM (Fig. 6E), indicating physical blockade of the access of Ubiquitin and NEM to E2-active site. These findings establish that zinc ions inhibit hUBE2O by sterically blocking its E2 catalytic center through coordination with specific cysteine residues in the UBC domain.

The combined structural and biochemical evidence reveal a precise molecular mechanism whereby zinc ions bind to cysteine residues in hUBE2O’s catalytic domain, physically obstructing substrate access to the active site and thereby inhibiting its enzymatic activity. This allosteric inhibition mode differs from the activating effect observed in E3-RING domains, highlighting enzyme-specific regulation by metal ions in the ubiquitination system.

## Discussion

Through a combination of domain truncation, systematic mutagenesis and well-designed biochemical strategies, our study establishes that E2-conjugating enzyme hUBE2O achieves E3-independent ubiquitination through a unique mechanistic paradigm characterized by structural domain cooperativity and structural flexibility. Unlike HECT and RBR E3-ligases, hUBE2O lacks a catalytic cysteine critical for E3 activity, ruling out ubiquitination mechanistic similarity to these E3 families. Although RING E3s require zinc ions for proper structural folding (9, 20), zinc ions inhibit hUBE2O’s activities, yet hUBE2O may share mechanistic features with RING E3s in Ubiquitin transfer. Previous studies indicate that while Ubiquitin transfer from E2∼Ubiquitin to an amine of a substrate can occur spontaneously, this process is highly inefficient, the addition of a RING E3 massively enhances this process (29–31). RING E3s primarily function to prime Ubiquitin in the E2∼Ubiquitin complex and juxtapose substrate lysine with the E2∼Ubiquitin thioester for efficiently Ubiquitin transfer. In our study, we show that the UBC domain alone cannot form the thioester intermediate with Ubiquitin or catalyze self-ubiquitination and polyubiquitin chain formation without the CTR domain (Fig. 3, B and E). These findings indicate that the CTR domain serves as a multifunctional module: facilitating both E2∼Ubiquitin intermediate formation and subsequent turnover to lysine residues of substrates. Furthermore, the structural flexibility of the CTR domain is essential for hUBE2O’s catalytic function (Fig. 3F), suggesting its conformational plasticity enables the allosteric changes required for efficient Ubiquitin transfer.

In addition to the UBC and CTR domains, the CC domain is critical for substrate ubiquitination, as evidenced by its critical role in ubiquitination of CTNNA1 (Fig. 4, C and D) and SMAD6 in previous study (14). Notably, residue T995, located proximal to the CC domain, influences substrate selectivity, implying the CC domain may facilitate proper positioning of lysine residues of substrates relative to the hUBE2O∼Ubiquitin thioester bond for efficient Ubiquitin transfer. Therefore, unlike classical E2 enzymes that rely on E3 ligases for substrate recruitment and Ubiquitin-ligating activity amplification, hUBE2O’s bifunctionality arises from cooperation between its UBC catalytic core and flanking CC/CTR domains. This domain architecture design allows hUBE2O to bypass E3 partners while maintaining efficient Ubiquitin transfer to lysine residues and substrate specificity, defining it as a hybrid enzyme with E2-E3 functional duality. Notably, zinc ions and phenylarsenoxides inhibit hUBE2O by crosslinking its C910 and C913 residues (Figs. 6B, S1C and S5C), implying that the conformational changes through these vicinal cysteines are required for catalysis.

Our identification of six mutations (T995A, S1042A, S1046A, S1060A, H939A, and H1130A) that impair hUBE2O activity (Figs. 4A and S5B) reveals critical regulatory nodes in its catalytic mechanism. All six mutations reduce E2∼Ubiquitin adduct formation, with H939A, T995A, and S1024A exhibiting particularly pronounced effects (Fig. 4E). AlphaFold3 modeling of hUBE2O-D2 (CC, UBC and CTR domains)-Ubiquitin complex (Fig. S3C) suggests that S1042 and S1060 are essential for efficient ubiquitination probably by mediating UBC domain and Ubiquitin interactions. While previous study established the functional importance of dimerization for the tpUBE2O activity, our data show that H939A, T995A, and H1130A mutants of hUBE2O retain dimerization capability (Fig. S5D), indicating activity defects of these three mutants likely stem from other mechanisms that warrant further investigation.

Counterintuitively, hUBE2O activity is resistant to its phosphorylation and self-ubiquitination, which are the modifications that typically fine-tune E2 and E3 function (9, 19, 24). Instead, the identification of zinc ions as potent allosteric inhibitors of hUBE2O unveils a novel regulatory axis in ubiquitination. Whereas zinc ions are essential for RING E3 domain folding and activity (9, 20), their inhibitory effect on hUBE2O highlights context-dependent metal-mediated regulation within ubiquitination system. Biochemical evidences and structural modeling reveal that zinc ions binding to cysteines (C910, C913 and C1020) induces steric occlusion of the catalytic pocket, blocking Ubiquitin access to E2 active site C1040. Importantly, zinc sensitivity is conserved in drUBE2O despite partial sequence divergence, implying evolutionary conservation of this regulatory mechanism in vertebrates. The differential zinc sensitivity between hUBE2O and other RING-E3s enzymes may reflect their distinct biological roles, with hUBE2O inhibition serving a safeguard against uncontrolled E3-independent ubiquitination. Moreover, the N-terminal CR1-CR2 regions of hUBE2O mediated interactions with substrates including SMAD6, RECQL4 and CTNNA1, establishing their autoinhibitory role (Fig. 3G) as an orthogonal regulatory mechanism that prevents non-specific ubiquitination events. We hypothesize that the substrate binding induces a conformational switch that relieves autoinhibition, enable hUBE2O to function as a potent E2/E3 enzyme capable of ubiquitinating these substrates.

Although AlphaFold3 predictions offer critical structural insights, experimental validation of hUBE2O’s conformational dynamics is essential to characterize its functional flexibility. Despite extensive efforts to determine the structure of hUBE2O under diverse crystallization conditions, the flexible nature of its CTR region precluded structural resolution. Additionally, no specific interacting partners were identified in our hands that could stabilize the CTR domain, further challenging structural characterization. The physiological relevance of zinc-mediated inhibition of hUBE2O enzymatic activity also warrants future structural and biological investigation. Furthermore, as an E2 enzyme with E3-mimetic activity, the functional crosstalk between hUBE2O and canonical E3 ligases represents an important avenue for further exploration.

The deletion variants and site-directed mutants generated in this study enable the generation of substrate proteins with defined ubiquitination signatures (varying in modification sites and chain topology), which can serve as powerful tools for dissecting Ubiquitin code-dependent cellular processes. Furthermore, hUBE2O has been implicated in paradoxical tumor-promoting and tumor-suppressive roles, underscoring its context-dependent biological functions (21, 22). Our mechanistic discoveries expand the therapeutic potential for targeting ubiquitination pathways, as hUBE2O’s non-catalytic domains, residues, and auto-inhibition represent novel druggable sites. For instance, the T995A mutation was identified in pancreatic ductal adenocarcinoma (COSMIC database: https://cancer.sanger.ac.uk), highlighting the clinical relevance of this regulatory hotspot. Moreover, the zinc ions-mediated allosteric regulation of hUBE2O presents a novel therapeutic paradigm for modulating its activity in disease contexts.

## Experimental procedures

### Plasmids, transfection and cell culture

The full-length and truncated versions of pCR3.1-hUBE2O-Myc, pCR3.1-hUBE2O-C1040S-mutated (CS)-Myc, pLV-Flag-hUBE2O-TS, pLV-Flag-hUBE2O and pLV-Flag-hUBE2O-C1040S-mutated (CS) plasmids have been described previously (3, 32). For the deleted versions of hUBE2O, the indicated regions were cloned into N-terminal Flag-tagged pLV vector. The single cysteine mutated hUBE2O plasmids were mutated based on pCR3.1-hUBE2O-Myc plasmid and the single or combined serine, tyrosine, threonine, histidine and lysine mutated plasmids were mutated based on pLV-Flag-hUBE2O plasmid using Phanta Max Super-Fidelity DNA Polymerase (Vazyme, P505-d2). The h*UBE2O-K0* (all lysine residues are replaced with arginine residues) fragment and zebrafish *Ube2o* fragment were synthesized from IGE Bio, China, and subsequently cloned into CMV promoter-based N-terminal Flag-or Myc-tagged vectors. The full-length pCR3.1-HA-mCTNNA1, pGEX-6P-1-mCTNNA1-Myc, pGEX-6P-1-mCTNNA1-V5 and pCR3.1-HA-RECQL4 plasmids have been described before (3). The full-length human *BMAL1* was amplified from ORF of CCSB-Broad Lentiviral Expression Library (Accession: BC041129) and cloned into N-terminal Flag-tagged pCR3.1 vector. The mUBE1, Ubiquitin WT and K0 plasmids have been described previously (3, 4). All plasmids were confirmed by DNA sequencing. The sequence of oligos used for plasmids construction are shown in Table S1. Poly-ethylenimine (PEI, Polysciences, 24765-1g) reagent was used for plasmids transfection.

Expi293F^TM^ cells were obtained from Gibco (A14527) and maintained in SMM 293-TII Expression Medium (Sino Biological, M293TII). HEK293T (CRL-3216^TM^) cells were purchased from ATCC. HEK293T cells were routinely maintained in Dulbecco’s Modified Eagle’ s Medium (DMEM, Gibco, C11995500BT) supplemented with 10% Fetal bovine serum (FBS, Gibco, 10270106) in a humidified atmosphere containing with 5% CO_2_ at 37°C. All cells were tested for mycoplasma once per month routinely using PCR based method (33).

#### Western blot and antibodies

Cells were lysed in lysis buffer (0.5% NP40, 150 mM NaCl, 50 mM Tris pH 8.0, 1mM EDTA, 10% Glycerol (Sigma-Aldrich, V900122)) supplemented with 1×cOmplete^TM^ Protease Inhibitor Cocktail (Roche, 4693116001) on ice for 30 min. The lysates were then centrifuged at 20,000 *g* for 10 min at 4°C. The supernatants were transferred into a new tube and the pellet were washed by PBS, resuspend and lysed in the same volume of lysis buffer containing 1% SDS and 1×Protease Inhibitor Cocktail at 95°C for 15 min. For the total extract, cells were lysed in lysis buffer supplemented with 1% SDS and 1×Protease Inhibitor Cocktail at 4°C for 30 min. The lysates were sonicated by Biorupter Pico with 10 cycles of 30 s on and off, and then heated at 95°C for 5 min. The protein concentrations of the lysates were measured using the Pierce^TM^ BCA Protein Assay Kit (ThermoFisher Scientific, 23225). Equivalent amounts of proteins were boiled with 1×LDS loading buffer for 5 min at 95°C, resolved by SDS–PAGE and transferred to nitrocellulose membrane (Cytiva, 10600003) for immunoblotting. Membranes were blocked with 5% non-fat milk (Beyotime, P0216) in TBST (50 mM Tris-HCl pH 7.6, 150 mM NaCl, 0.1% Tween 80) and incubated with indicated primary antibodies overnight at 4°C. The membranes were washed 3 times with TBST by the adding of goat anti-rabbit (Bethyl Laboratories, A120-101P; 1:10,000), or goat anti-mouse (Bethyl Laboratories, A90-116P; 1:10,000) secondary HRP-conjugated antibody for 1 h at room temperature. After 3 times washes, images were taken using ChemiDoc Imaging systems from Bio-Rad using BeyoECL Star substrate (Beyotime, P0018AS).

Primary antibodies used in western blot included rabbit anti-Myc (Proteintech, 16286-1-AP; 1:5,000), rabbit anti-HA (Sigma-Aldrich, H6908; 1:5,000), mouse anti-Flag tag (Sigma-Aldrich, F1804; 1:5,000), rabbit anti-Flag (Merck millipore, F7425; 1:1,500; Figs. 3B, 3D-F and 6A), rabbit anti-UBE2O (Cell signaling technology, 83393S; 1:2,000), rabbit anti-V5 tag (GeneTex, GTX117997; 1:5,000), mouse anti-Phospho-threonine (42H4) (Cell signaling technology, 9386, 1:1,000), mouse anti-α-Tubulin (DM1A) (Abcam, ab7291; 1:4,000), mouse anti-ubiquitin (P4D1) (Santa Cruz, SC-8017, 1:1000) and mouse anti-GAPDH (Proteintech, 60004-1-Ig; 1:10,000). For Coomassie blue staining, Imperial Protein Stain (ThermoFisher Scientific, 24615) was used.

### Protein purification and *in vitro* ubiquitination assay

His-HA-Ubiquitin-WT, His-HA-Ubiquitin-K0, Flag-UBE2O-TwinStrep, CTNNA1-Myc, CTNNA1-V5, His-mUBE1, UBE2D1, GST-MARCH7 RING purified proteins were from our previous works (4). For in vitro ubiquitination assay of purified hUBE2O protein, 2 μΜ Flag-UBE2O-TwinStrep, 25 μM His-HA-ubiquitin, 100 nM His-mUBE1 with or without 2 μΜ mCTNNA1-Myc were incubated at 37°C for 1 h in ubiquitination reaction buffer containing 2 mM ATP, 5 mM MgCl_2_, 50 mM NaCl, and 50 mM Tris-HCl pH 8.0. The reaction was quenched with 1×LDS loading buffer for 5 min at 95°C.

For in vitro ubiquitination assay of hUBE2O deletions and site-specific mutations, HEK293T cells transfected with the indicated plasmids for 36 h were harvested and lysed by lysis buffer supplemented with 1×Protease Inhibitor Cocktail. The lysates were centrifuged at 20,000 *g* for 10 min at 4°C. A portion of the supernatants were collected as input samples, while the remaining supernatants were subsequently incubated with 5 μL of pre-washed anti-Flag M2 affinity gel (Sigma-Aldrich, A2220) for 60 min at 4°C, or 30 μL of Pierce™ Anti-c-Myc Magnetic Beads (ThermoFisher Scientific, 88843) for 90 min at 4°C, followed by 3 times washes with lysis buffer. Subsequently, *in vitro* ubiquitination reactions were performed by incubating beads at 37°C for 1 h with 25 μM His-HA-Ubiquitin and 100 nM mE1 in ubiquitination reaction buffer with or without 2 μΜ CTNNA1-Myc/V5 in a table shaker. Proteins bound to beads were eluted and the reactions were quenched with 1×LDS loading buffer for 15 min at 95°C.

### In vitro dephosphorylation followed by ubiquitination assay

HEK293T cells transfected with Flag-hUBE2O-WT or Flag-hUBE2O-C1040S-mutated plasmid for 36 h were harvested and lysed by lysis buffer supplemented with 1×Protease Inhibitor Cocktail. The lysates were centrifuged at 20,000 *g* for 10 min at 4°C. A portion of the supernatants were collected as input samples, while the remaining supernatants, or purified Flag-hUBE2O-TwinStrep protein, were incubated with anti-Flag M2 affinity gel for 60 min at 4°C followed by 3 times washes with lysis buffer. The beads were then incubated with or without Lambda Protein Phosphatase (Lambda PP, New England Biolabs, P0753S), FastAP™ Thermosensitive Alkaline Phosphatase (Thermo Fisher Scientific, EF0651) or Quick CIP (New England Biolabs, M0525S) according to manufacturer’s protocol in a table shaker, followed by 3 times washes with lysis buffer to remove the phosphatase. Subsequently, in vitro ubiquitination reactions were performed by incubating beads at 37°C for 1 h with 25 μM His-HA-Ubiquitin and 100 nM mE1 in ubiquitination reaction buffer in a table shaker. Proteins bound to beads were eluted and the reactions were quenched with 1×LDS loading buffer for 15 min at 95°C.

### E2∼Ubiquitin adducts formation and turnover assay

For in vitro E2∼Ubiquitin adducts formation assay of purified hUBE2O protein, 0.4 μΜ Flag-hUBE2O-TwinStrep, 25 μM His-HA-ubiquitin, 100 nM mE1 were incubated at room temperature for 3 min after mixing with ubiquitination reaction buffer. The reaction was quenched with 1×LDS loading buffer with or without 2-Mercaptoethanol (Sigma-Aldrich, M3148) for 5 min at 95°C.

For in vitro E2∼Ubiquitin adduct formation assay of hUBE2O deletions and site-specific mutations, HEK293T cells transfected with the indicated plasmids for 36 h were harvested and lysed by lysis buffer supplemented with 1×Protease Inhibitor Cocktail. The lysates were centrifuged at 20,000 *g* for 10 min at 4°C. A portion of the supernatants were collected as input samples, while the remaining supernatants were subsequently incubated with 6 μL pre-washed anti-Flag M2 affinity gel (Sigma-Aldrich, A2220) for 60 min at 4°C followed by 3 times washes with lysis buffer. Subsequently, beads were incubated at room temperature for 1 min or 3 min (as indicated in the figures) after mixing with 25 μM His-HA-Ubiquitin and 100 nM mE1 in ubiquitination reaction buffer in a table shaker. The reaction was quenched with 1×LDS loading buffer with or without 2-Mercaptoethanol for 5 min at 95°C.

For in vitro E2∼Ubiquitin adducts turnover assay of purified hUBE2O protein (Fig. 2, B and C), 70 μL of ubiquitination reaction containing 0.4 μM Flag-hUBE2O-TwinStrep, 25 μM His-HA-ubiquitin, 100 nM mE1 and ubiquitination reaction buffer were mixed on ice. A 9 μL aliquot of the reaction was mixed with 1 μL of 0.8 mM N-Ethylmaleimide (NEM, Sigma-Aldrich, E3876) (sample 3#). After 3 min of room temperature incubation, the reaction was quenched by putting on ice rapidly. Two 9 μL aliquots of the reaction were mixed with 1 μL of 0.5 M EDTA on ice (samples 4# and 6#), two 9 μL aliquots of the reaction were mixed with 1 μL of 0.8 mM NEM on ice (samples 5# and 7#) and two 9 μL aliquots of the reaction were mixed with 1 μL of 0.5 M EDTA and 10 μL of 4×LDS loading buffer with (sample 2#) or without (sample 1#) 2-Mercaptoethanol. Samples 3-7# were further incubated at 37°C for 1 min (Samples 4# and 5#) or 5 min (Samples 6# and 7#), followed quenching through adding of 10 μL of 4×LDS loading buffer. Samples 1-7# were boiled at 95°C for 5 min before separated by SDS–PAGE and analyzed by immunoblotting.

### Immunoprecipitation assay

Cells transfected with the indicated plasmids for 36 h were washed twice with ice-cold PBS and lysed by lysis buffer supplemented with 1×Protease Inhibitor Cocktail. The lysates were centrifuged at 20,000 *g* for 10 min at 4°C. A portion of the supernatants were collected as input samples, while the remaining supernatants were subsequently incubated with pre-washed anti-Flag M2 affinity gel for 90 min at 4°C followed by 3 times washes with lysis buffer. Bound proteins were eluted by boilling in 2×LDS loading buffer at 95°C for 15 min. The input samples and eluates were separated by SDS–PAGE and analyzed by immunoblotting.

### Ubiquitination assay in HEK293T cells

Ubiquitination assay in HEK293T cells by nickel pull-down (PD: Ni-NTA) was performed as previously described (14). Briefly, HEK293T cells transfected with indicated plasmids were washed twice with ice-cold PBS containing 10 mM NEM. Cells were lysed in a denaturing buffer (8 M urea, 0.1 M NaH_2_PO_4_-Na_2_HPO_4_, 10 mM Tris-HCl pH 7.0, 10 mM imidazole) supplemented with 10 mM 2-Mercaptoethanol for 15 min at room temperature. The lysates were clarified by centrifugation at 20,000 *g* for 10 min at room temperature. A portion of the supernatants were collected as input samples, while the remaining supernatants were subsequently incubated with pre-washed nickel agarose (QIAGEN, 30230) at room temperature for 2 h with gentle rotation. Beads were washed 3 times with denaturing buffer and bound proteins were eluted in 2×LDS loading buffer at 42°C for 15 min. The input samples and eluates were separated by SDS–PAGE and analyzed by immunoblotting.

Ubiquitination assay in HEK293T cells by anti-Flag immunoprecipitation (IP: Flag) was conducted as previously described (3). In brief, HEK293T cells transfected with indicated plasmids for 36 h were washed twice with ice-cold PBS containing 10 mM NEM. The cells were lysed in radioimmunoassay buffer (20 mM NaH_2_PO_4_-Na_2_HPO_4_ pH 7.4, 150 mM NaCl, 1% Triton X-100, and 0.5% sodium deoxycholate) supplemented with 1% SDS, 1×Protease Inhibitor Cocktail and 10 mM NEM at 4°C for 30 min. The lysates were sonicated by Biorupter Pico with 10 cycles of 30 s on and off, and then heated at 95°C for 5 min. Subsequently, the cell lysates were diluted to 0.1% SDS by radioimmunoassay buffer and clarified by centrifugation at 20,000 *g* at 4°C for 10 min. A portion of the supernatants were collected as input samples, while the remaining supernatants were subsequently incubated with pre-washed anti-Flag M2 affinity gel for 90 min at 4°C, followed by 3 times washes with radioimmunoassay buffer. The bound proteins were eluted in 2×LDS loading buffer at 42°C for 15 min. The input samples and eluates were separated by SDS–PAGE and analyzed by immunoblotting.

### PAO affection on ubiquitination assay

For PAO affection assay of hUBE2O cysteine-to-serine mutations in Fig. S1C, HEK293T cells transfected with the indicated plasmids for 36 h were harvested and lysed by lysis buffer supplemented with 1×Protease Inhibitor Cocktail. The lysates were centrifuged at 20,000 *g* for 10 min at 4°C. A portion of the supernatants were collected as input samples, while the remaining supernatants were subsequently incubated with pre-washed anti-c-Myc magnetic beads for 90 min at 4°C, followed by 3 times washes with lysis buffer. Subsequently, beads were incubated with 0.08 mM PAO (Sigma-Aldrich, P3075) in buffer containing 50 mM NaCl, 25 mM Tris pH 7.5 and 10% Glycerol at 37°C for 20 min. Beads were washed 3 times with lysis buffer and incubated with 25 μM His-HA-Ubiquitin and 100 nM mE1 in ubiquitination reaction buffer in a table shaker at 37°C for 1h. The reaction was quenched with 1×LDS loading buffer at 95°C for 15 min.

### Metal ion affection on ubiquitination assay

For metal ion affection on ubiquitination assay of Fig. 5C, 0.4 μΜ purified Flag-UBE2O-TwinStrep protein was incubated with 0.1 mM indicated metal ions in ubiquitination reaction buffer with or without ATP at 37°C for 1 h. 25 μM His-HA-ubiquitin, 100 nM mE1 and 2 μΜ CTNNA1-Myc were added for another 1 h incubation at 37°C. The reaction was quenched with 1×LDS loading buffer for 5 min at 95°C.

For zinc affection assay of hUBE2O WT, deletions, site-specific mutations and zebrafish homolog, HEK293T cells transfected with the indicated plasmids for 36 h were harvested and lysed by lysis buffer supplemented with 1×Protease Inhibitor Cocktail. The lysates were centrifuged at 20,000 *g* for 10 min at 4°C. A portion of the supernatants were collected as input samples, while the remaining supernatants were subsequently incubated with pre-washed anti-Flag M2 affinity gel for 60 min at 4°C, or anti-c-Myc magnetic beads for 90 min at 4°C, followed by 3 times washes with lysis buffer. Subsequently, beads were incubated with zinc chloride in Tris buffer (50 mM Tris pH 7.5 and 200 mM NaCl) at the indicated concentration at 37°C for 1 h. Beads were washed 3 times with lysis buffer and incubated at 37°C for 1h with 25 μM His-HA-Ubiquitin and 100 nM mE1 in ubiquitination reaction buffer with or without 2 μΜ CTNNA1-Myc in a table shaker. The reaction was quenched with 1×LDS loading buffer at 95°C for 15 min.

For zinc specificity assay of Fig. 5, D and E, 1.3 μΜ purified GST-MARCH7 RING or Flag-UBE2O-TS was incubated with 0.1 mM zinc chloride in buffer containing 50 mM NaCl, 25 mM Tris pH 7.5 and 10% Glycerol at 37°C for 1 h. The reaction was clarified by centrifugation at 20,000 *g* at 4°C for 10 min. A portion of the supernatants were collected as input samples, while the remaining supernatants were subsequently diluted 20-fold in lysis buffer and incubated with pre-washed anti-Flag M2 affinity gel for 60 min at 4°C, followed by 3 times washes with lysis buffer. Beads was incubated with RING ubiquitination reaction mixture (containing 25 μM His-HA-ubiquitin, 100 nM mE1, 1 μM UBE2D1 and ubiquitination reaction buffer) or hUBE2O ubiquitination reaction mixture (containing 25 μM His-HA-ubiquitin, 100 nM mE1, 2 μM CTNNA1-Myc and ubiquitination reaction buffer) at 37°C for 1 h in a table shaker. The reaction was quenched with 1×LDS loading buffer at 95°C for 15 min.

### Zinc Colorimetric Assay

The concentration of free zinc ions in solution was quantified using a Zinc Colorimetric Assay Kit (Elabscience, E-BC-K137-M) according to the manufacturer’s protocol. Briefly, 2 μM purified Flag-UBE2O-TwinStrep protein was incubated with or without 0.1 mM zinc chloride in Tris buffer at 37°C for 1 h. The reaction mixture was then diluted 8-fold with ultra-pure water and mixed with the protein precipitator. After centrifugation at 13,780 *g* for 10 min at 4°C, the supernatant was collected and reacted with chromogenic working solution. The absorbance at 560 nm was measured after 30 s of shaking followed by 5 min incubation at room temperature. Zinc ion concentrations were calculated based on a standard curve generated with known concentrations of zinc standards.

### SEC-MALS

100 μL of 2 mg/mL Purified Flag-UBE2O-TwinStrep was loaded onto a Superdex 200 Increase 5/150 GL column (Cytiva, 28-9909-45) with buffer containing 50 mM Tris-HCl pH7.4, 200 mM NaCl, 0.5 mM TCEP. A static light-scattering detector and a differential refractive index detector (Wyatt Technology) were connected to the analytical gel filtration chromatography system. Data analysis was performed using ASTRA 6.1.1.17 software provided by Wyatt Technology.

### Proteomics sample preparation and mass spectrometry analysis

For identification of hUBE2O-catalyzed self-ubiquitination sites and polyubiquitin chain types, 1 μΜ Flag-UBE2O-TwinStrep and 100 nM mE1 were incubated with or without 25 μM His-HA-Ubiquitin WT or K0 at 37°C for 1 h in ubiquitination reaction buffer. The reaction was quenched with 1×LDS loading buffer for 5 min at 95°C. The reaction mixtures were separated by SDS–PAGE and stained by Coomassie Blue. The indicated bands were excised for in-gel digestion with trypsin and mass spectrometry analysis. 2-chloroacetamide instead of iodoacetamide was used for alkylation for ubiquitination sites identification. For mass spectrometry identification, tryptic peptides were separated using 140 min of total data collection time (100 min of 2% to 22%, 20 min of 22% to 28%, 12 min of 28% to 36%, 2 min of 36% to 100% and 6 min of 100% of buffer B (80% acetonitrile (ThermoFisher Scientific, 51101) in 0.1% formic acid (v/v) in water (ThermoFisher Scientific, 85170)) with a 300 nL/min flow using an Easy-nLC 1200 connected online to a Fusion Lumos (with FAIMS pro) mass spectrometer (ThermoFisher Scientific). Scans were collected in data-dependent top-speed mode with dynamic exclusion at 60 s.

For NEM-modified C1040 peptides identification, three repeats were performed. Specifically, 1.3 μΜ purified Flag-UBE2O-TS was incubated with 0.25 mM zinc chloride in Tris buffer at 37°C for 1 h in a table shaker. 20 μM NEM was subsequently added to the reaction and incubated at 37°C for 15 min in a table shaker. The reaction was clarified by centrifugation at 20,000 *g* at 4°C for 10 min. A portion of the supernatants were collected as input samples, while the remaining supernatant was diluted 20-fold in lysis buffer (no EDTA) and incubated with pre-washed anti-Flag M2 affinity gel for 60 min at 4°C, followed by 3 times washes with lysis buffer (no EDTA) and once wash with PBS. Proteins bound to beads were subjected to on-beads digestion with trypsin as described before (34). For mass spectrometry identification, tryptic peptides were separated using 60 min of total data collection time (2 min of 4% to 8%, 43 min of 8% to 28%, 8 min of 28% to 36%, 2 min of 36% to 100% and 5 min of 100% of buffer B in 0.1% formic acid (v/v) in water with a 300 nL/min flow using an Easy-nLC 1200 connected online to a Fusion Lumos mass spectrometer (ThermoFisher Scientific). Scans were collected in data-dependent top-speed mode with dynamic exclusion at 60 s.

Raw data were analyzed using MaxQuant (version 2.4.2.0) searched against hUBE2O FASTA (Q9C0C9) alone or together with human Ubiquitin FASTA (P62987, P62979, P0CG48 and P0CG47), with label-free quantification, match between runs functions (35), the GlyGly or NEM remnants search enabled. The output protein lists are available in datasets S1 and visualized using DEP, ggplot2, tidyr, dplyr, ggpubr, readr and trackViewer packages (36). The accession number for the raw mass spectrometry data reported in this paper is ProteomeXchange: PXD064901, now accessible at https://www.iprox.cn/page/PSV023.html;?url=1749735964100rQd2, password: mnld.

### Multiple sequence alignment of UBE2Os across different species

UBE2O protein sequences from different species including *Homo sapiens*, *Mus musculus*, *Xenopus laevis*, *Danio rerio*, *Drosophila rhopaloa*, and *Trametes pubescens* are downloaded from UniProt (https://www.uniprot.org/) with UniProtKB entries of Q9C0C9, Q6ZPJ3, A0A1L8EMX4, A0A8M3B1J4, A0A6P4EMW5 and A0A1M2VY70, respectively. Alignment was conducted with online tools of Clustal Omega (37) and ESPRIPT (38). Sequence numbering and secondary structural elements are shown according to the chain A of the complex of hUBE2O with NAP1L1 structure (PDB: 7UN6).

### Protein structure prediction using AlphaFold3

The structural model of the hUBE2O-D2/Ubiquitin complex was generated using AlphaFold3 (28) web server (https://alphafoldserver.com/). One copy of the hUBE2O D2 (801-1292aa) protein sequence and one copy of Ubiquitin protein sequence were input into the AlphaFold3 modeling pipeline to predict protein structures with high accuracy. Five model predictions were generated, S1042 and S1060 interact with Ubiquitin in all the five predicted models, and the one with high confidence score was selected for visualization.

The structural model of the hUBE2O and six zinc ions complex was generated using AlphaFold3 web server (https://alphafoldserver.com/). One copy of the hUBE2O full length protein sequence together with six zinc ions were input into the AlphaFold3 modeling pipeline to predict protein structures with high accuracy. Five model predictions were generated, and the one with the interaction between C910/C913 residues and zinc, together with the wild-type hUBE2O structure downloaded from AlphaFold Protein Structure Database (https://www.alphafold.ebi.ac.uk/) were selected for the overall structure and catalytic core analysis.

### Image analyses and quantifications

Analyses of the integrated density of the western blot images in Figs. 2C and 5F were performed using a built-in Fiji (39) macro, version 2.9.0. The “Measure” tool was used after subtracting background and inverting.

### Statistical analyses

Statistical evaluation of the data was performed using GraphPad Prism (version 10.2.3). All quantifications represent the mean ± SEM. Comparisons were carried out by One-way ANOVA followed by Tukey’s post hoc test. The differences were considered to be statistically significant at p<0.05 in the analytical treatment of the data.

## Data availability

The data underlying Figs. 1B, 1C, 6E and S2A have been deposited to the ProteomeXchange Consortium (https://proteomecentral.proteomexchange.org) via the iProX partner repository (40) with the dataset identifier PXD064901, now accessible at https://www.iprox.cn/page/PSV023.html;?url=1749735964100rQd2, password: mnld. Other data are available in the article itself and its supporting information.

## Supporting information

This article contains supporting information. There are 5 supplemental figures, 1 supplemental table, 1 supplemental dataset and 11 blots SourceDatas associated with this work.

## Acknowledgments

We thank all members of Zhang lab for their constructive suggestions and helps. This work was financially supported by grants from National Natural Science Foundation (32370752), Guangdong Basic and Applied Basic Research Project (2024A1515012344), and the Science and Technology Planning Project of Guangdong Province (2023B1212060050, 2023B1212120009).

## Author Contributions

D.X. and X.Z. designed the experiments. X.T. and R.S. performed the mass spectrometry measurement. D.X. and X.T. analyzed the mass spectrometry data. S.D. performed the SEC-MALS analysis. D.X. performed all the remaining experiments. D.X. and X.Z. wrote the manuscript.

## Conflict of interest

The authors declare that they have no conflicts of interest with the contents of this article.

### Abbreviations

The abbreviations used are: UBE2O: (E3-independent) ubiquitin-conjugating enzyme E2O;
PTMs: Post-translational modifications;
E1: ubiquitin-activating enzymes;
E2: ubiquitin-conjugating enzymes;
E3: ubiquitin ligases;
RING: really interesting new gene;
HECT: homologous to E6AP C-terminus;
RBR: RING-between-RING;
CR: conserved region;
CC: coiled coil;
UBC: ubiquitin conjugating,
CTR: C-terminal regulatory;
K0: lysine-deficient or lysine-null, all lysine residues are replaced with arginine residues;
SEC-MALS: Size Exclusion Chromatography–Multi-Angle Light Scattering;
NEM: N-Ethylmaleimide;
Cys^E3^: a E3 catalytic cysteine.
MLL: mixed-lineage leukemia;
Lambda PP: Lambda Protein Phosphatase;
FastAP: FastAP Thermosensitive Alkaline Phosphatas;
IB: immunoblotting;
CB: Coomassie blue staining;
FL: full-length;
WT: wild-type,
LFQ: label free quantification
2-Me: 2-Mercaptoethanol.

---

**Figure S1.**
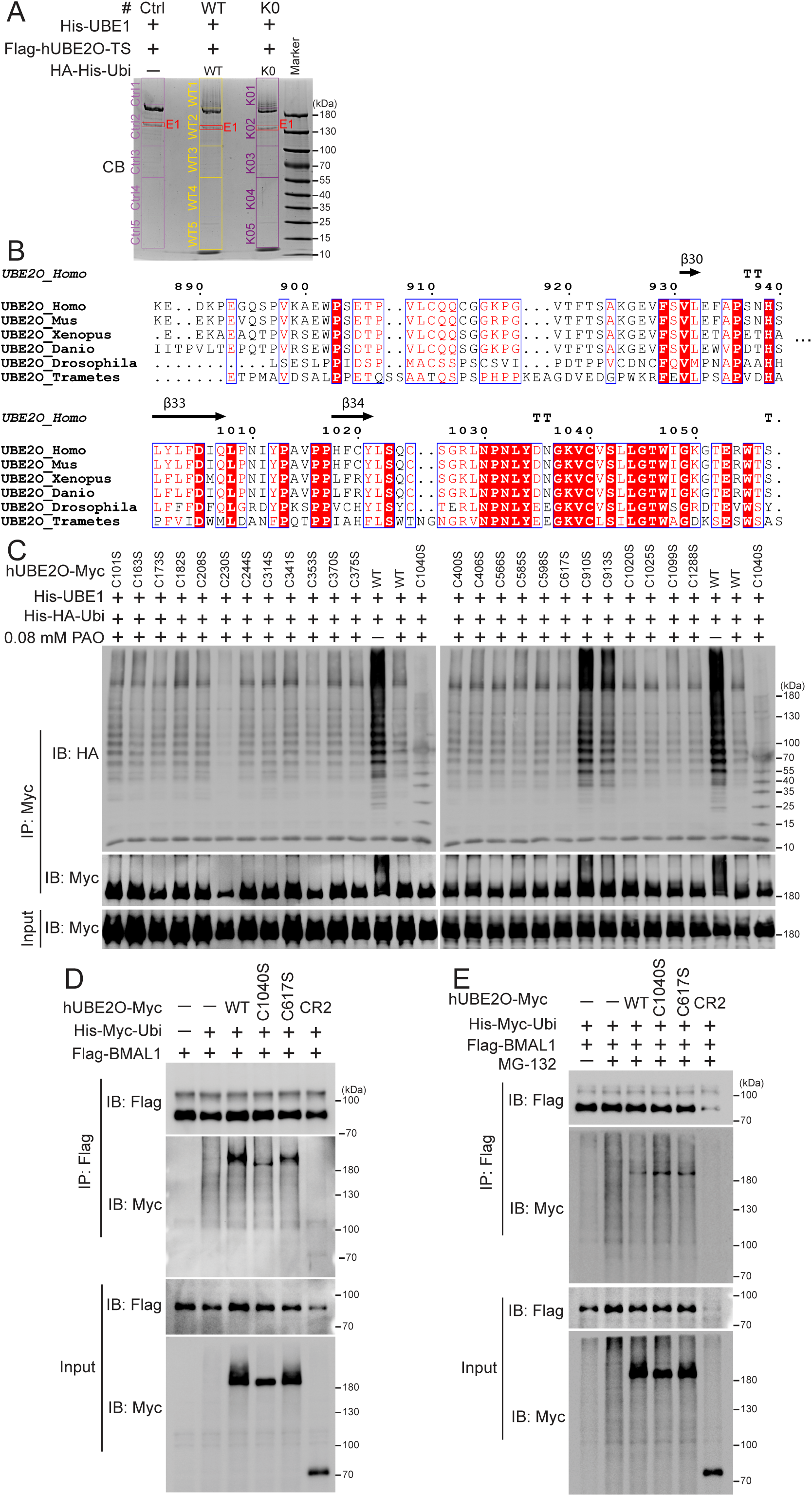
C1040 is the sole catalytic cysteine of hUBE2O. *A,* Coomassie Blue staining of the indicated ubiquitination reactions separated by SDS– PAGE. The indicated bands (excluding E1) were excised and subjected to in-gel tryptic digestion followed by mass spectrometry analysis. *B,* multiple sequence alignment of UBE2Os across different species. Identical residues are colored in red and specially highlighted with red frames; similar residues across the group are highlighted in blue boxes and colored in red, with exceptions in black. *C,* immunoblots show C910S and C913S mutations are resistant to PAO-mediated inhibition among all 25 hUBE2O cysteine to serine mutations. *D* and *E*, ubiquitination assays in HEK293T cells show neither full-length hUBE2O nor C617 mutant catalyze detectable BMAL1 ubiquitination with (*E*) or without (*D*) 12 h treatment of 10 mM MG-132 under our experimental conditions. Experiments were repeated at least twice, one representative result is shown. Source data for this figure are available in supporting information as: SourceDataFS1.

**Figure S2.**
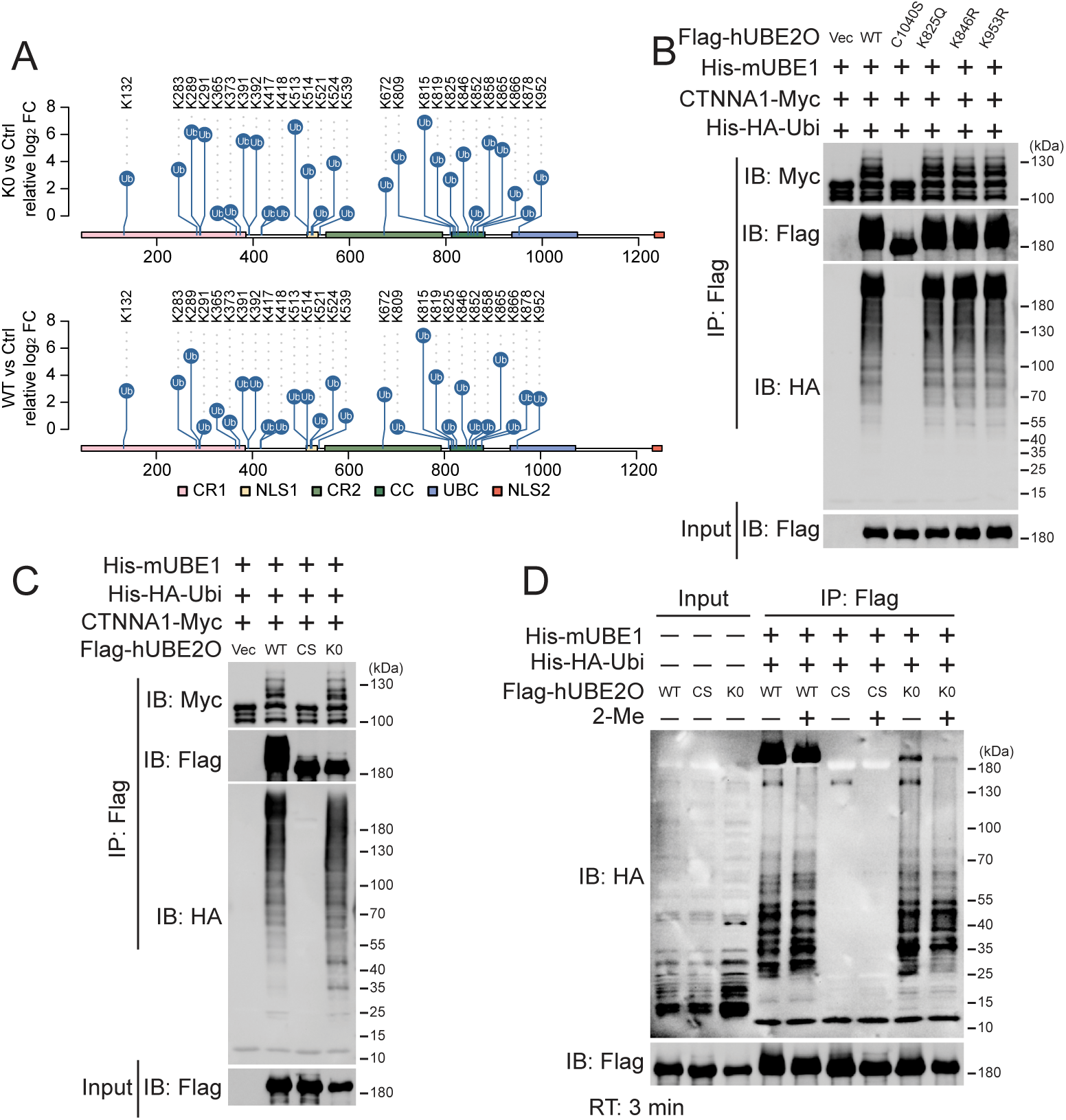
Self-ubiquitination barely regulates hUBE2O activity. *A,* schematic presentation of the location of hUBE2O self-ubiquitinylated sites detected by mass spectrometry. Detailed data are available in datasets S1. K0: lysine-null. *B,* in vitro ubiquitination assay shows the self-ubiquitination (IB: Flag), polyubiquitin chain formation (IB: HA) and CTNNA1 ubiquitination catalyzing (IB: Myc) ability of the indicated hUBE2O WT, C1040S and KR mutations expressed in HEK293T cells. *C,* in vitro ubiquitination assay shows the self-ubiquitination (IB: Flag), polyubiquitin chain formation (IB: HA) and CTNNA1 ubiquitination catalyzing (IB: Myc) abilities of the indicated hUBE2O WT, C1040S (CS) and lysine-null (K0) mutations expressed in HEK293T cells. *D,* E2∼Ubiquitin adducts formation assay shows the E2∼Ubiquitin adducts and polyubiquitin chain formation abilities of the indicated hUBE2O WT, C1040S (CS) and lysine-null (K0) mutations expressed in HEK293T cells. All experiments except for mass spectrometry identification were repeated at least twice, one representative result is shown. Source data for this figure are available in supporting information as: SourceDataFS2.

**Figure S3.**
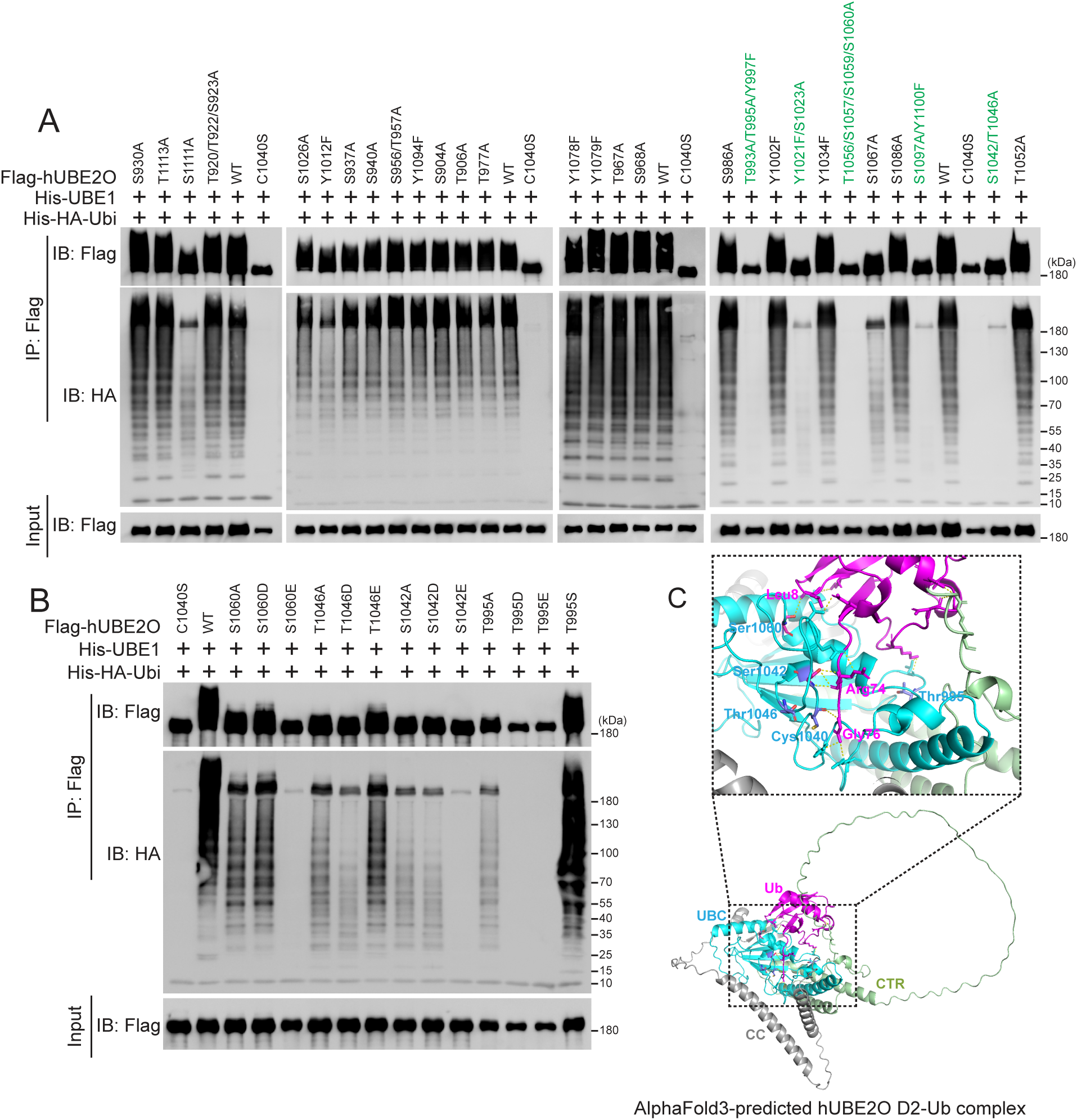
hUBE2O’s activity is resistant to phosphorylation. *A,* in vitro ubiquitination assay shows the self-ubiquitination (IB: Flag) and polyubiquitin chain formation (IB: HA) abilities of the HEK293T-expressed hUBE2O WT, C1040S and mutations of indicated serine, threonine and tyrosine residues in the UBC domain. *B,* in vitro ubiquitination assay shows the self-ubiquitination (IB: Flag) and polyubiquitin chain formation (IB: HA) abilities of the HEK293T-expressed hUBE2O WT, C1040S and the phosphorylation-prevented or phosphorylation-mimetic mutations of S1060, T1046, S1042 and T995 residues. *C,* a structural model of hUBE2O-D2 (801-1292aa) in complex with Ubiquitin generated using AlphaFold3. hUBE2O is shown as cartoon representation with individual domains labeled and color coded. The CC domain of hUBE2O (gray), UBC domain of hUBE2O (cyan), the CTR domain of hUBE2O (green), and Ubiquitin (magenta) are shown as cartoons (bottom). The inter-domain interaction residues of hUBE2O are shown as cyan sticks (S1060, T1046, S1042, C1040 and T995 residues are highlighted as slate sticks), nitrogen and oxygen atoms are respectively in blue and red), and the corresponding interaction residues of Ubiquitin are shown as magenta sticks. All experiments were repeated at least twice, one representative result is shown. Source data for this figure are available in supporting information as: SourceDataFS3.

**Figure S4.**
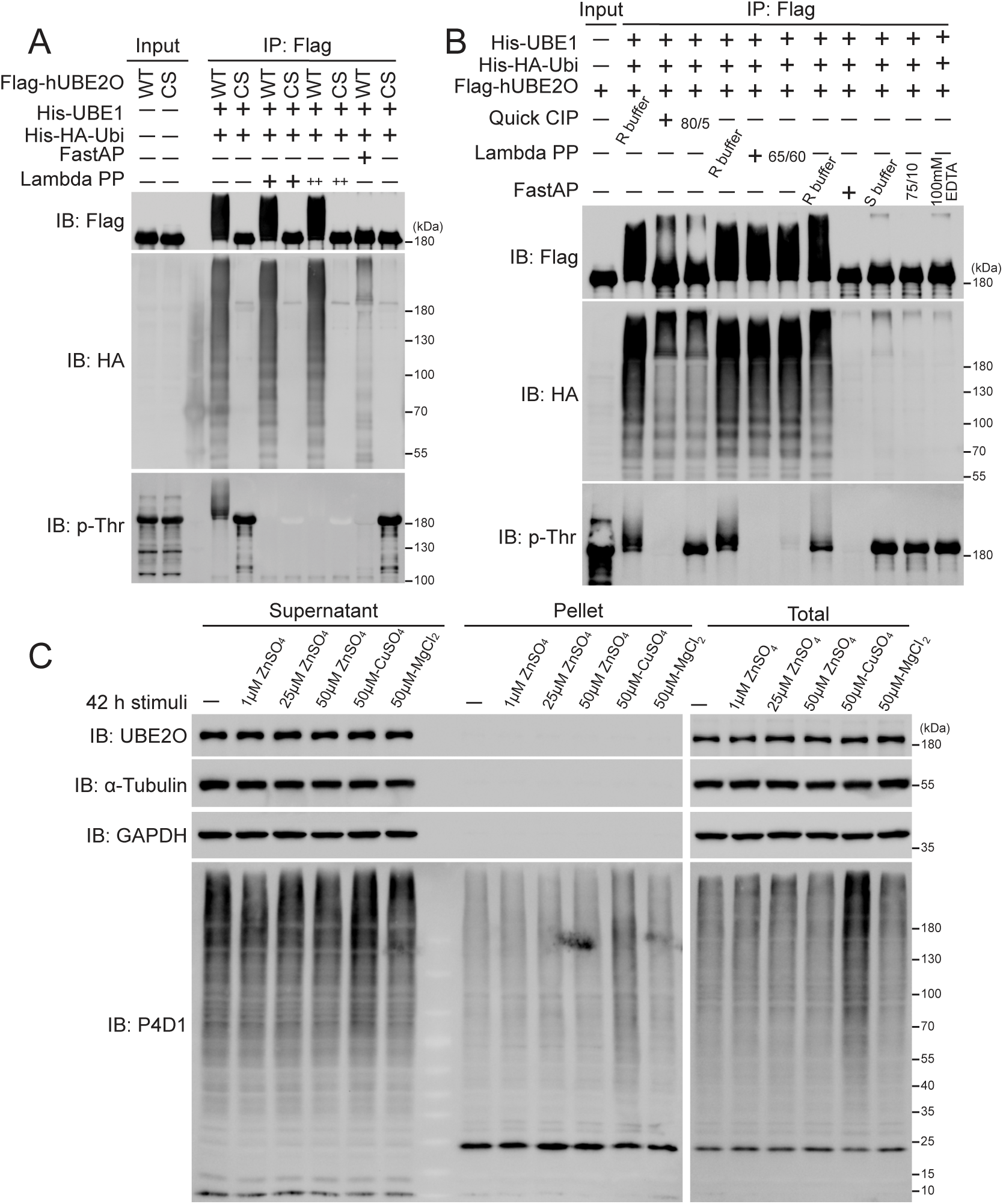
Zinc ions, but not phosphorylation inhibits the enzymatic activity of hUBE2O. *A,* in vitro dephosphorylation followed by ubiquitination assay shows Lambda PP and FastAP treatment successfully reduce threonine phosphorylation of hUBE2O (IB: p-Thr) but only FastAP treatment completely abolishes hUBE2O’s self-ubiquitination (IB: Flag). ++ indicates 2 ×Lambda PP phosphatase was used. *B,* in vitro dephosphorylation followed by ubiquitination assay shows the storage buffer of FastAP, but not dephosphorylation abolishes hUBE2O’s self-ubiquitination. Quick CIP, Lambda PP and FastAP were inactivated by incubating at 80°C for 5 min (80/5), 65°C for 60 min (65/60) and 75°C for 10 min (75/10), respectively. FastAP was also inactivated by 100 mM EDTA before dephosphorylation assay. *C,* immunoblots show zinc ions barely affect endogenous UBE2O solubility/accumulation or overall polyubiquitination patterns in HEK293T cells. HEK293T cells treated with the indicated metal ions for 48 h were collected for western blot analysis according to the experimental procedures. P4D1 antibody was used to detect ubiquitin, polyubiquitin and ubiquitinated proteins. All experiments were repeated at least twice, one representative result is shown. Source data for this figure are available in supporting information as: SourceDataFS4.

**Figure S5.**
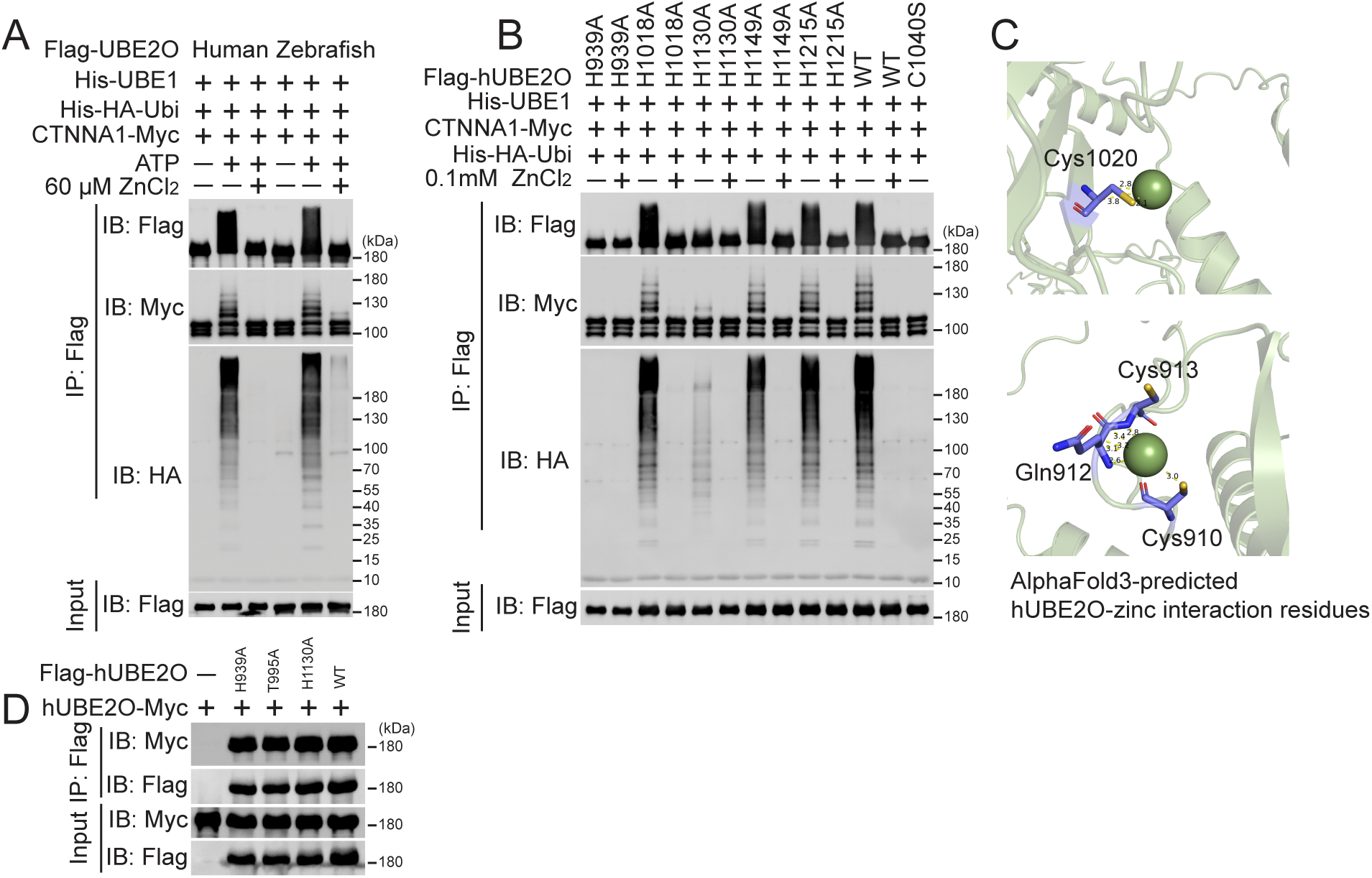
Zinc ions coordinate with specific cysteine residues in hUBE2O. *A* and *B,* in vitro ubiquitination assay shows the self-ubiquitination (IB: Flag), polyubiquitin chain formation (IB: HA) and CTNNA1 ubiquitination catalyzing (IB: Myc) abilities of the indicated hUBE2O, its zebrafish homolog and histidine to alanine mutations expressed in HEK293T cells. *C,* structural models of the interaction residues of hUBE2O to zinc ions in the hUBE2O-6zinc complex generated using AlphaFold3. hUBE2O is shown as green cartoon. The interaction residues of hUBE2O are shown as slate sticks, nitrogen, oxygen and sulfur atoms are respectively in blue, red and yellow. *D,* immunoprecipitation assay shows the interactions between hUBE2O WT and the H939A, T995A or H1130A mutations. All experiments were repeated at least twice, one representative result is shown. Source data for this figure are available in supporting information as: SourceDataFS5.

**Table S1.**
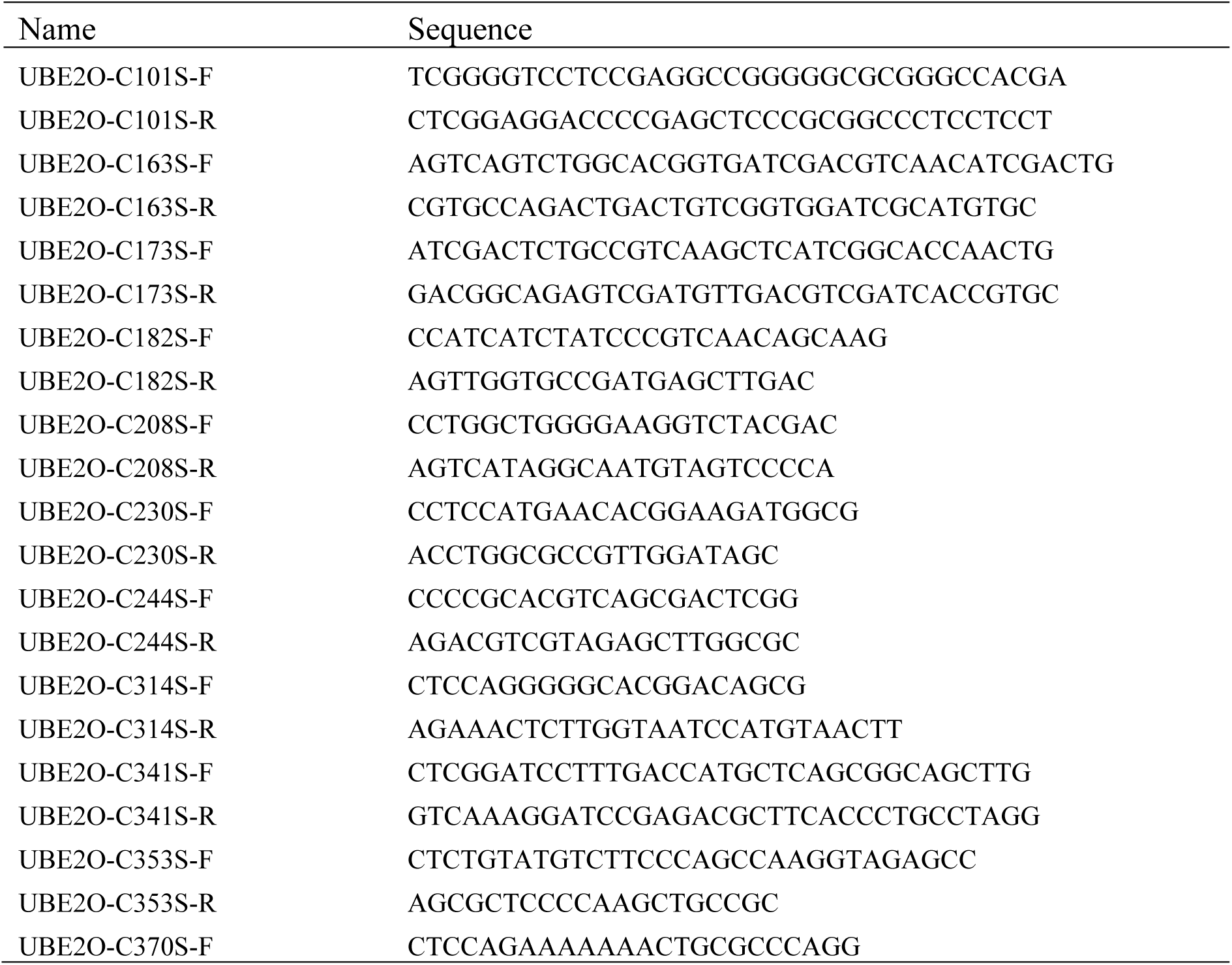
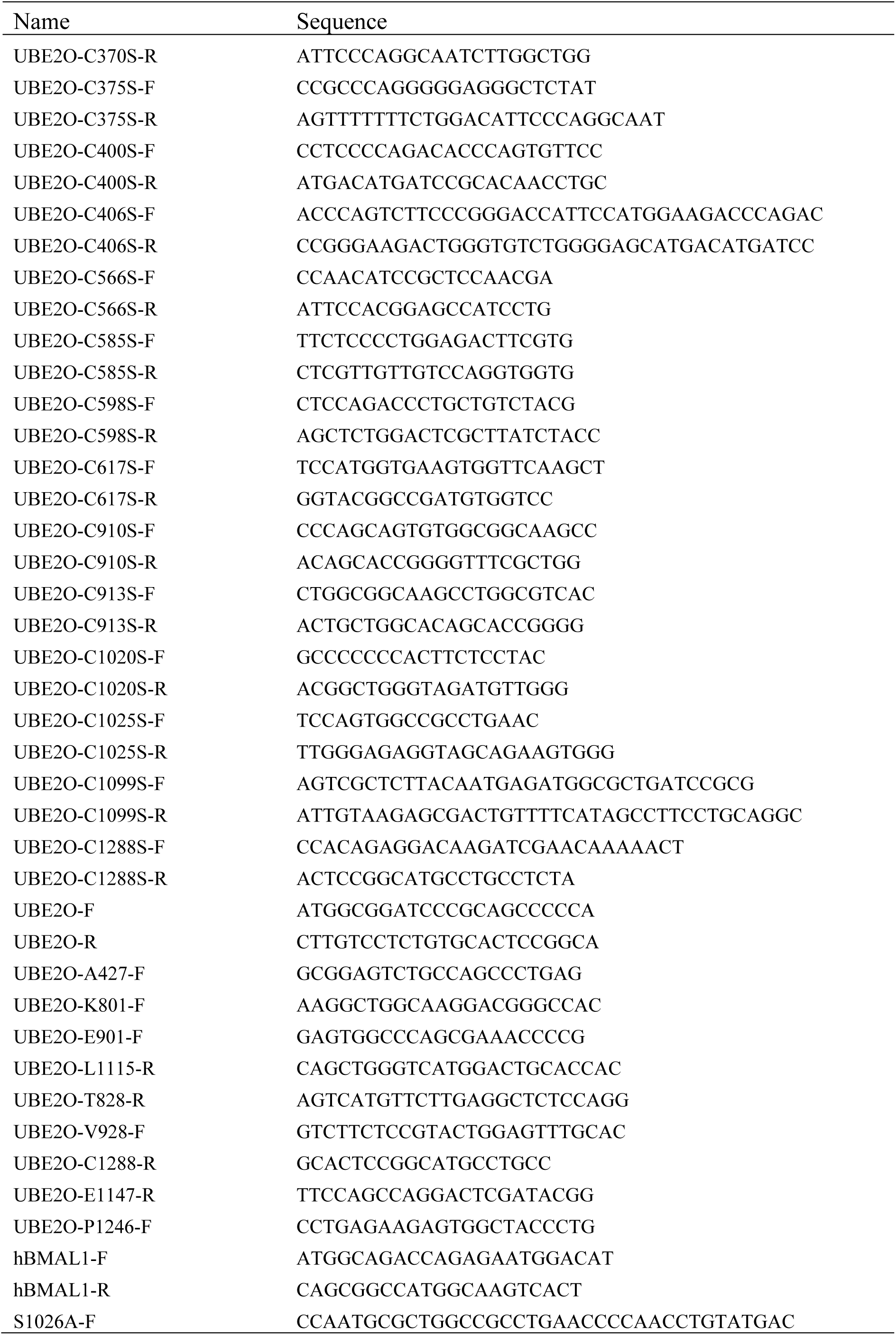
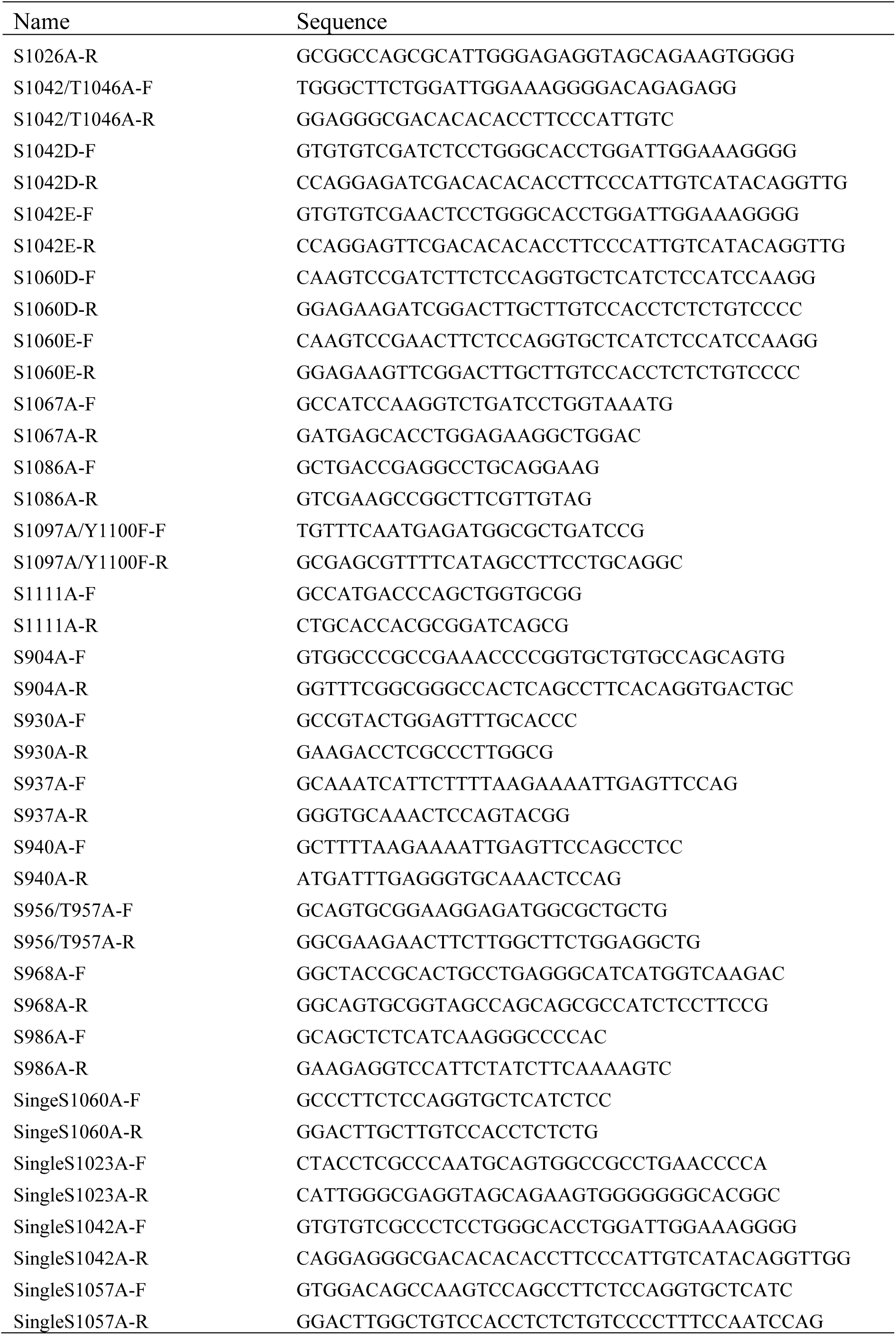
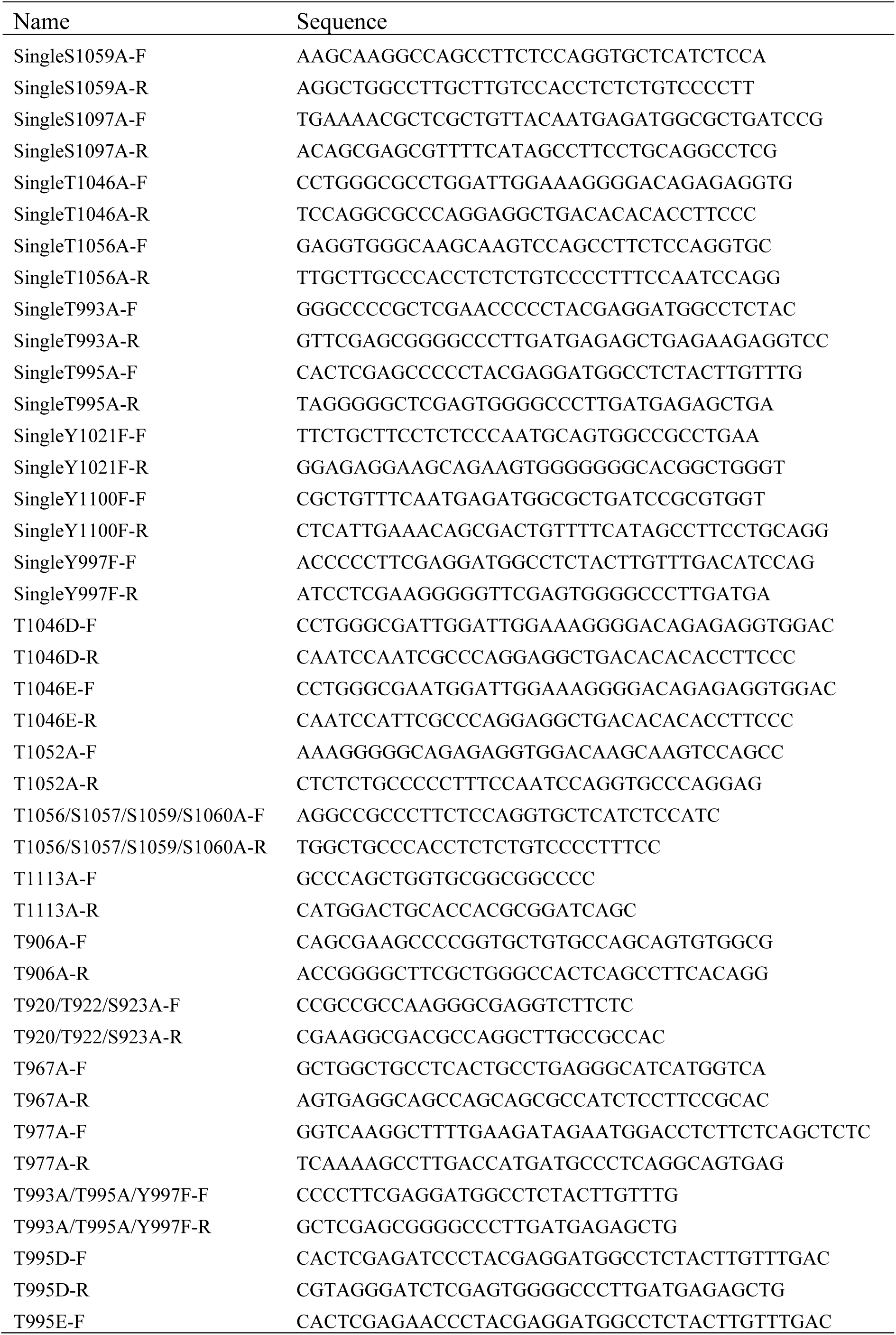
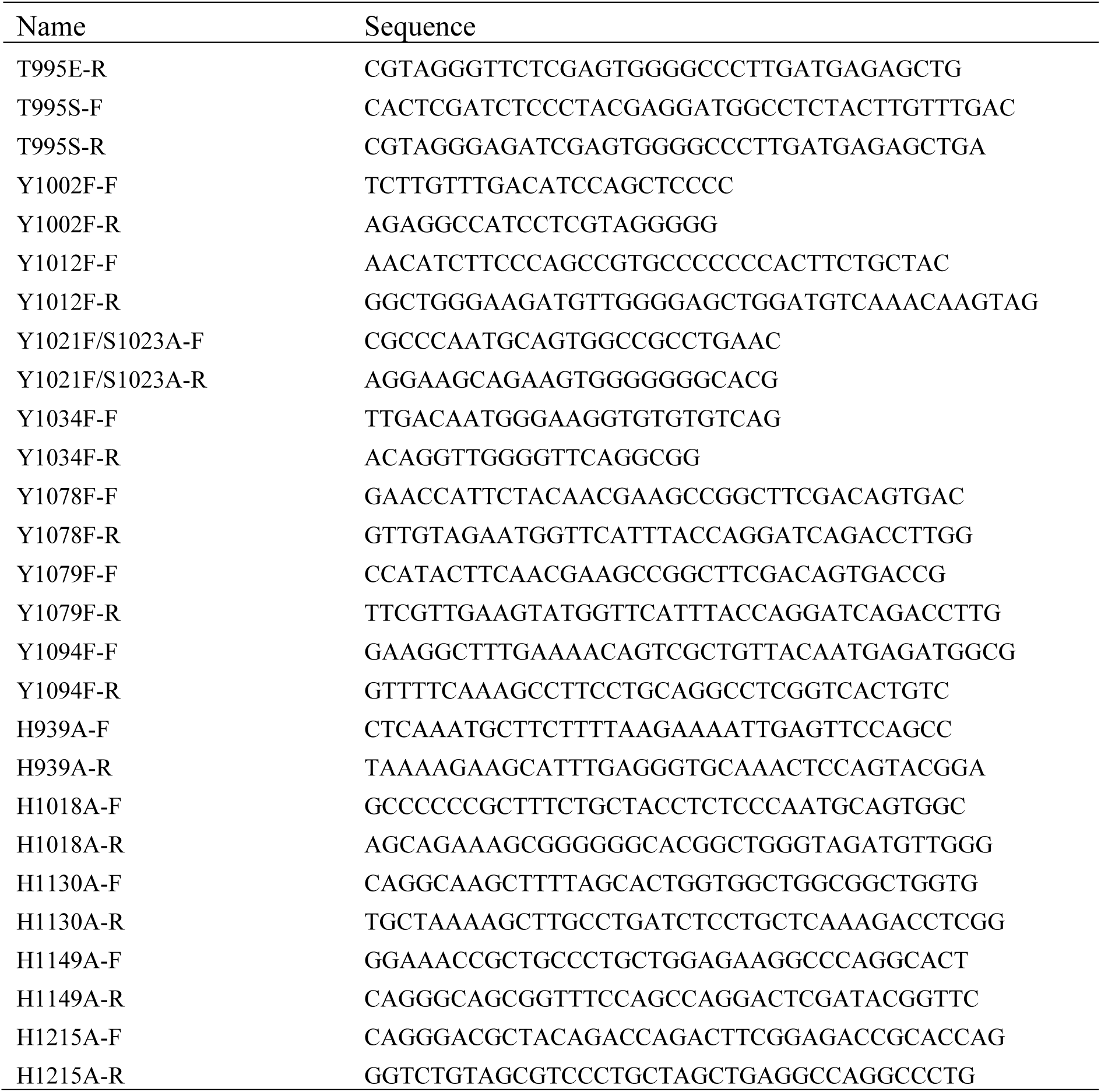
Oligos used in this study.

